# Reduced glucosinolate content in oilseed rape (*Brassica napus* L.) by random mutagenesis of *BnMYB28* and *BnCYP79F1* genes

**DOI:** 10.1101/2022.08.10.503450

**Authors:** Srijan Jhingan, Hans-Joachim Harloff, Amine Abbadi, Claudia Welsch, Martina Blümel, Deniz Tasdemir, Christian Jung

## Abstract

The presence of anti-nutritive compounds like glucosinolates (GSLs) in the rapeseed meal severely restricts its utilization as animal feed. Therefore, reducing the GSL content to <18 µmol/g dry weight in the seeds is a major breeding target. While candidate genes involved in the biosynthesis of GSLs have been described in rapeseed, comprehensive functional analyses are missing. By knocking out the aliphatic GSL biosynthesis genes *BnMYB28* and *BnCYP79F1* encoding an R2R3 MYB transcription factor and a cytochrome P450 enzyme, respectively, we aimed to reduce the seed GSL content in rapeseed. After expression analyses on single paralogs, we used an ethyl methanesulfonate (EMS) treated population of the inbred winter rapeseed ‘Express617’ to detect functional mutations in the two gene families. Our results provide the first functional analysis by knock-out for the two GSL biosynthesis genes in winter rapeseed. We demonstrate that independent knock-out mutants of the two genes possessed significantly reduced seed aliphatic GSLs, primarily progoitrin. Compared to the wildtype Express617 control plants (36.3 µmol/g DW), progoitrin levels were decreased by 55.3% and 32.4% in functional mutants of *BnMYB28* (16.20 µmol/g DW) and *BnCYP79F1* (24.5 µmol/g DW), respectively. Our study provides a strong basis for breeding rapeseed with improved meal quality in the future.

## Introduction

Oilseed rape or rapeseed (*Brassica napus* L.) is an essential oil crop, ranking as the third-largest source of vegetable oil globally (http://www.fao.org/faostat/). In Europe, it is grown as a winter crop, sown in autumn and flowering in the following spring after exposure to cold temperatures over winter. The seeds contain 45-50% oil with a healthy lipid profile, suitable for human consumption ^[1]^. Moreover, it is utilized for biodiesel production. After oil extraction, the rapeseed meal (RSM) serves as a protein-rich (40%) animal feed. However, major anti-nutritive compounds like glucosinolates (GSLs) in RSM adversely affect its nutritional and commercial value ^[2]^. Therefore, increasing yield potential, seed oil content, and improving seed meal quality are major goals for rapeseed breeding.

GSLs are diverse heterogeneous secondary metabolites specific to Brassicales. They are sulfur and nitrogen-containing products derived from glucose and amino acids as precursors, comprising a thioglucose and a sulfonated oxime attached to the chain elongated amino acid. Depending on their respective amino acid precursors, GSLs are categorized as aliphatic, aromatic, and indolic, originating primarily from methionine, phenylalanine, and tryptophan, respectively ^[3]^. Biosynthesis of the three types is independently controlled by distinct gene families ^[4]^. Roughly 130 different GSL types have been described from 16 dicot angiosperms^[5]^, most of which are edible plants ^[6; 7]^. Apart from *B. napus, B. oleracea* (cauliflower, cabbage, broccoli, Brussels sprouts, and kale), and *B. rapa* (turnips and radish) are economically relevant vegetables rich in GSLs ^[8; 4]^. Fifteen major GSL types have been identified in *B. napus* ^[6]^, reaching levels as high as 60-100 µmol/g dry weight in seeds with the methionine-derived aliphatic GSLs constituting up to 92% of all GSL types ^[9]^. The development of rapeseed varieties with low GSL seed content was a milestone in rapeseed breeding ^[10]^. Alleles conferring low seed GSL content were introgressed from the Polish spring variety ‘Bronowski’ ^[11]^ to develop modern rapeseed cultivars with improved seed meal traits. In modern varieties, the GSL content of the RSM has been reduced to 8-15 μmol per gram seed weight ^[10]^.

GSL biosynthesis is completed in three major steps, i) chain elongation, ii) core structure formation, and iii) secondary side-chain modifications ^[3]^. First, the addition of methylene groups results in chain elongated amino acids. Next, the addition of the sulfur group to the chain-elongated amino acids and S-glucosylation completes the core structure formation. Lastly, secondary modifications like benzoylation, desaturation, hydroxylation, methoxylation, and oxidation result in distinct GSL types ^[4; 7]^. Environmental effects combined with specific genetic mechanisms for GSL biosynthesis, regulation, transport, and storage result in varying GSL contents and diverse profiles observed across *Brassica* species^7^

GSLs yield toxic by-products after enzymatic cleavage by the endogenous thioglucoside glucohydrolase called myrosinase ^[3]^. Upon physical injury, the myrosinase is released from so-called ‘myrosin cells’ and comes into contact with GSLs stored in ‘S-cells’ ^[12]^. Hydrolysis of GSLs generates various products like isothiocyanates (ITC), thiocyanates (SCN), epithionitriles, and nitriles (NI), many of which are known to confer defense against generalist herbivores and bacterial and fungal pathogens ^[13]^. GSLs have been shown to confer antimicrobial properties against the phytopathogenic bacterium *Xanthomonas campestris* pv. *campestris* and the necrotrophic fungus *Sclerotinia sclerotiorum*^[14]^. High consumption of GSLs through the feed can result in several adverse metabolic effects in animals.

Hydroxyalkenyl GSLs like epiprogoitrin and progoitrin are goitrogenic by causing inflammation of the thyroid gland ^[6]^. Retarded growth, reduced appetite, and feed efficiency, gastrointestinal irritation, liver and kidney damage, and behavioral effects have been observed in fish ^[15]^, poultry ^[16]^, and higher mammals like pigs ^[2]^. On the contrary, other GSL types like sulforaphane and indole-3-carbinol are known for their beneficial effects on human health with anti-carcinogenic properties ^[17]^.

A complex network of transcription factors (TFs) influenced by abiotic and biotic stimuli, hormonal and epigenetic factors controls the spatiotemporal regulation of GSL biosynthesis ^[18; 19]^. The most notable genes controlling aliphatic GSL biosynthesis in *Arabidopsis* are R2R3 MYB transcription factors. Three TFs *MYB28, MYB76*, and *MYB29*, also referred to as *HIGH ALIPHATIC GLUCOSINOLATE* (*HAG*) *1, 2*, and *3*, respectively, have been described ^[18; 19]^. *HAG1* has been speculated to have a ‘master regulator’ effect by up-regulating almost all genes involved in the core structure formation of aliphatic GSLs ^[19]^. In *Brassica* field crops, the role of *MYB28* genes have been demonstrated to be strongly associated with aliphatic GSL biosynthesis in *B. oleraceae* ^[20]^, *B. juncea* ^[21; 22; *23]*^ *and B. rapa* ^[24]^.

The gene *CYP79F1* controls the first step of the core structure formation by converting chain elongated methionine to corresponding aldoximes ^[25; 26]^. Its role in GSL biosynthesis has also been demonstrated in *B. juncea* ^[27]^ and *B. oleraceae* ^[28]^.

The transfer of knowledge from *Arabidopsis* to *B. napus* is limited and complicated due to its polyploid genome, where multiple genes with functional redundancies may exist. Several genes associated with GSLs in rapeseed have been revealed through associative transcriptomics, genome-wide association, and QTL mapping studies ^[29; 30; 31; 32; 33]^. These studies demonstrated the significant association between biosynthesis genes *BnMYB28* and *BnCYP79F1* with high aliphatic GSL content.

Our study aimed to reduce aliphatic GSLs in rapeseed since they are the most abundant in seeds. We analyzed the expression profiles of three *BnMYB28* and two *BnCYP79F1* paralogs in rapeseed. Then, we selected *BnMYB28*.*C09, BnMYB28*.*A03, BnCYP79F1*.*C05* and *BnCYP79F1*.*A06* as candidate genes for functional studies. Using an ethyl methanesulfonate (EMS) mutagenized winter rapeseed population ^[34]^, we detected loss-of-function mutations in *BnMYB28* and *BnCYP79F1* genes involved in the core structure biosynthesis of aliphatic GSLs. Double mutants displayed a significant reduction in the seed aliphatic GSL content.

These materials could be interesting for breeding rapeseed with improved seed meal quality by achieving a further reduction of aliphatic GSLs in the seeds.

## Materials and Methods

### Plant material and growth conditions

We used the oilseed rape EMS population previously developed using an advanced inbred line (F_11_) of the winter rapeseed variety ‘Express’ ^[34]^. Seeds were treated with 0.5-1.2% EMS for 12 h. The resulting M_2_ plants were selfed to produce the corresponding M_3_ populations.

M_3_ plants were grown in 11 cm pots under greenhouse conditions (16 h light, 20-25°C) for three weeks, with non-mutagenized plants of Express617 as controls. They were vernalized for eight weeks in a cold chamber (16 h light, 4°C). Plants selected for crossing experiments were hand-pollinated after emasculation. The inflorescences of plants chosen for selfings were isolated with plastic bags before anthesis. Plants selected for GSL measurements were grown in 11 cm pots under greenhouse conditions (16 h light, 20-25°C).

### DNA isolation and PCR

For genomic DNA isolation, leaf samples were collected and lyophilized for 72 h (Martin Christ Gefriertrocknungsanlagen GmbH, Germany). Freeze-dried samples were pulverized using the GenoGrinder2010 (SPEX^®^ SamplePrep LLC, USA) at 1,200 strokes/min. Genomic DNA was isolated using the standard CTAB method ^[35]^. PCR was performed using paralog-specific primers as per the following conditions: 94°C for 2 min, 36 cycles of 94°C for 30s, 58-66°C for 30 s - 1 min and 72°C for 1 min, followed by 72°C for 5 min for final elongation.

### Bioinformatics analyses

Genomic DNA and polypeptide sequences of aliphatic GSL biosynthesis genes *AtMYB28* and *AtCYP79F1* were retrieved from The Arabidopsis Information Resource (TAIR - https://www.arabidopsis.org/). Using the Darmor-*bzh* rapeseed reference genome (http://www.genoscope.cns.fr/brassicanapus/), amino acid sequences of the *Arabidopsis* orthologs were used as BLAST queries. Retrieved chromosomal locations and gene sequences of hits with the lowest e-values and >80% sequence similarity were accepted. Gene annotations (exons, 5’ and 3’ untranslated regions and open reading frames) for the paralogs were made using the CLC Main workbench 7 (QIAGEN^®^ Aarhus A/S, Aarhus C, Denmark). Sequence alignments were generated for the genomic DNA, cDNA, and amino acid sequences of the retrieved genes. Conserved and functional domain analyses were done using the NCBI Conserved Domain Database.

### Gene expression analysis by RT-qPCR

The winter-type rapeseed inbred line Express617 was used for expression studies. Seeds were sown in 11 cm pots under greenhouse conditions (16 h light, ∼25°C). After three weeks, plants were vernalized (16 h light, 4°C) for eight weeks and then transferred to greenhouse conditions (16 h light, ∼25°C) and their positions were randomized twice a week. Flowers were hand-pollinated and marked with pollination dates. Leaves and seeds were sampled at 15, 25, 35, and 45 days after pollination (DAP). 50-100 mg of fresh weight tissues were collected from five biological replicates at the four developmental stages. Tissues were frozen in liquid nitrogen and stored at -70°C. Frozen tissues were pulverized in 2 ml reaction tubes with three 3 mm steel balls using the GenoGrinder2010 (SPEX^®^ SamplePrep LLC, USA) at 1,200 strokes/min in 1 min intervals. RNA was isolated using the peqGold Plant RNA Kit (PEQLAB Biotechnologie GmbH, Germany) following the manufacturer’s instructions. RNA quality was assessed with a NanoDrop2000 spectrophotometer (ThermoFisher Scientific, USA) and by agarose gel electrophoresis (1.5% agarose, 100 V, 30 min). The RNase-free DNase kit (ThermoFisher Scientific, USA) was used to treat samples with DNaseI to remove contaminating gDNA. cDNA was synthesized with 1 µg RNA using the First Strand cDNA Kit (ThermoFisher Scientific, USA). RT-qPCR was performed on the Bio-Rad CFX96 Real-Time System (Bio-Rad Laboratories GmbH, Germany) using paralog-specific primers (Supplementary Table 1). The relative expression was calculated according to the ΔΔC_q_ method for each paralog normalized against the two reference genes *BnGAPDH* and *BnACTIN*. The relative expression levels of each candidate paralogs were determined as a mean of five biological replicates with three technical replicates each.

### Conventional gel-based detection of EMS induced mutations

We screened 3,840 M_2_ plants from the EMS-mutagenized winter rapeseed Express617 population ^[34]^ to detect EMS-induced mutations in *BnMYB28* and *BnCYP79F1* paralogs. Paralog-specific primers were designed for the selected paralogs using the Darmor-*bzh* reference genome (Supplementary Table 2). Amplicons were evaluated for specificity using agarose gel electrophoresis (1% agarose, 100V, 10 min) and Sanger sequencing for validation. Using the protocol described by Till et al. (2006), we amplified M_2_ DNA pools using 5’ end infrared labeled probes DY-681 and DY-781 (Biomers, Ulm, Germany) for forward and reverse primers (100 pmol/µl), respectively. The resulting amplicons were processed for heteroduplex formation prior treatment with CELI nuclease (15 min at 45°C). 5 µl of 50 mM EDTA (pH 8.0) was added to terminate the digestion reaction. After digestion, samples were purified on Sephadex G-50 Fine columns (GE Healthcare, USA). 4 µl of formamide-containing dye (96% deionized formamide, 5 ml 0.25 M EDTA, 0.01% bromophenol blue) was added to each sample. Samples were concentrated to ∼20% of the original volume after incubation at 95°C for 30 min. 0.65 µl concentrated samples were separated on polyacrylamide gels using the LI-COR 4300 DNA Analyzer (LI-COR Biosciences, USA) using standard parameters (1,500 V, 40 mA, and 40 W for 4 h 15 min). The GelBuddy imaging software ^[37]^ was used to analyze gel images and identify single M_2_ mutants. Standard PCR was done to amplify regions harboring the expected single mutations using the gDNA isolated from single M_2_ plants. Amplicons were Sanger sequenced to validate the detected EMS-induced point mutations. Mutation effects conferred by SNPs on the polypeptide level were then characterized. Mutation frequencies (F) were estimated following the formula given by Harloff et al. (2012):

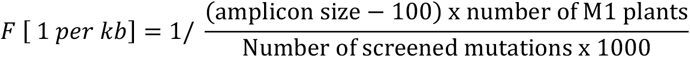

### Mutant genotyping

Using standard PCR, primers flanking detected EMS mutations were used for amplification (Supplementary Table 2). PCR specificity was checked using agarose gel electrophoresis (1%, 100V, 12-30 min). Plants were genotyped by Sanger sequencing of PCR fragments to confirm the presence of EMS mutations.

### Glucosinolate measurements

GSLs in leaves and mature seeds were measured in two ways. Quantitative measurements were performed with an enzymatic assay using myrosinase/thioglucosidase from *Sinapis alba* (Sigma-Aldrich CAS-No. 9025-38-1) and a D-Glucose Assay Kit (glucose oxidase/peroxidase; GOPOD assay) (Megazyme International, Ireland). A qualitative assessment of GSL profiles was done using high-performance liquid chromatography (HPLC). 5-7g fresh leaves (∼15 days after pollination) and 200-600 mg mature seeds were sampled in 50 ml and 2 ml sample tubes, respectively. Sampled leaves were lyophilized for 72 h (Martin Christ Gefriertrocknungsanlagen GmbH, Germany). Freeze-dried leaf samples and mature seeds (BBCH 89) were pulverized using the GenoGrinder2010 (SPEX^®^ SamplePrep LLC, USA) at 1,400 strokes/min in 4-5 1 min intervals. Using the hot methanol (70%) extraction method, ∼200 mg milled samples were used to prepare crude extracts following the protocol of Fiebig and Arens (1992) and then stored at -20°C.

For quantitative measurements, crude extracts (4 ml) were passed through 0.5 ml DEAE-Sephadex A-25 columns (GE Healthcare, USA). The bound GSL was digested on the column for 18 h at room temperature with ∼0.8U myrosinase/thioglucosidase. After digestion, columns were washed twice with 0.5 ml deionized distilled water (ddH_2_O) for elution of glucose. The eluate was shock frozen and freeze-dried for ∼72 h. The residue was dissolved in 100 µl ddH_2_O and a 5-40 µl aliquot was used for analysis in duplicates using the D-Glucose Assay Kit (Megazyme International, Ireland) following the manufacturer’s instructions.

For qualitative analyses, GSL in 1 ml crude extracts were bound on 250 µl DEAE-Sephadex A-25 columns and digested on the column for 18 h at room temperature with 18U sulfatase H1 enzyme (Merck KGaA, Germany). Desulfoglucosinolates were eluted twice with 1 ml ddH_2_O. 10-50 µl aliquots were separated on a 250 × 4.6 mm Lichrosorb 5µ column (Merck KGaA, Germany) as described by Fiebig and Arens (1992) using a Shimadzu2000 HPLC-system. GSLs were quantified by their absorbance at 229 nm and identified by retention time using commercially available GSL standards (PhytoLab GmbH, Germany). For each commercially available standard, an individual calibration curve was used for quantification (Supplementary Table 3).

To validate peak identity and quantification, a test set of 8 seed extract samples (concentration 8 mg/ml) from HPLC measurements were analyzed for cross-referencing on an LC-MS system consisting of a VWR Hitachi Elite LaChrom (Hitachi High-Technologies Corporation, Japan), L-2450 DAD detector, L-2300 Column Oven, L-2200 Autosampler and L-2130 Pump connected to an iontrap esquire4000 (Bruker Daltonics, Germany). The chromatography was carried out on a Synergi 4 µ Polar-RP column (80 Å, 250 × 4.6 mm, Phenomenex, Torrance, USA). The mobile phase consisted of H_2_O (Milli-Q grade, Arium^®^ Water Purification Systems, Sartorius, Germany) with 0.1% formic acid (Promochem, ScienTest-BioKEMIX GmbH, Germany) as eluent A, and acetonitrile (LC-MS-grade, AppliChem, Germany) as eluent B ^[39]^. The following chromatographic conditions were applied: elution starting from 0% B 0-2 min isocratic, gradient elution from 2-12 min to 60% B, 12-14 min isocratic elution at 60% B, gradient elution from 14-15 min to 0% B, 15-23 min isocratic re-equilibration at 0% B, 1 ml/min flow, oven temperature 30°C, wavelength 229 nm, injection volume 50 µl. The iontrap settings were as follows: negative mode, capillary voltage: 4000 V, nebulizer: 50 psi, dry gas (nitrogen): 10.0 L/min, dry temperature: 365°C, scan range: 100-1200 m/z ^[39]^.

Major sample peaks were quantified at 229 nm using a VWR Hitachi Chromaster (Hitachi High-Technologies Corporation, Japan) consisting of a 5430 DAD detector, 5310 Column Oven, 5260 Autosampler, and a 5110 Pump. The chromatographic conditions were applied as described for the LC-MS system, except that no formic acid was added to eluent A. The injection volume was 30 µl.

### Statistical analyses

For expression studies, significantly expressed paralogs were identified by performing an ANOVA (*p* <0.05) for the relative expression levels of each paralog. Mean relative expressions from five biological replicates were compared across the four sampling points 15, 25, 35, and 45 DAP in seeds and leaves. The LSD test (α ≤ 0.05) was performed to generate statistical groups using the ‘Agricolae’ package in R. The standard error of the mean was calculated across the five biological replicates with three technical replicates each.

In GSL determination experiments, the total GSL content using the D-Glucose Assay Kit and the contents of individual GSL compounds using HPLC were evaluated for statistical significance across the analyzed samples. An ANOVA (*p* <0.05) was performed for the analysis of the variance along with an LSD test (α ≤ 0.05) for statistical grouping using the ‘Agricolae’ package in R. The standard error of the mean was calculated across five biological replicates.

## Results

### Identification of *MYB28* and *CYP79F1* genes in the oilseed rape genome

For identification of possible paralogs in rapeseed, the Darmor-*bzh* reference genome (https://www.genoscope.cns.fr/brassicanapus/) was searched for *MYB28* and *CYP79F1* genes using polypeptide sequences from *A. thaliana* genes *AtMYB28* (AT5G61420) and *AtCYP79F1* (AT1G16410) as queries. Based on the lowest e-values and highest sequence similarities (>80%), three and two paralogs were detected for *BnMYB28* and *BnCYP79F1*, respectively (Table 1, Supplementary Figure 1). The polypeptides of *BnMYB28* and *BnCYP79F1* shared a similarity of 85% and 86%, respectively with their corresponding *Arabidopsis* orthologs. We aligned the polypeptide sequences of the candidate paralogs to identify conserved functional domains characteristic for the R2R3 MYB transcription factor and cytochrome P450 gene families (Supplementary Figure 2). In line with previous reports, the highly conserved DNA binding R2 and R3 domains specific to the subgroup 12 MYB transcription factors and the nuclear localization signal with the ‘LKKRL’ amino acid residues were present across protein sequences of all three *BnMYB28* paralogs ^[40]^. The paralog *BnMYB28*.*C09* was annotated in a truncated form in the Darmor-*bzh* reference genome but harbored all conserved domains required for gene activity. The protein sequences of both *BnCYP79F1* paralogs possessed five conserved domains characteristic to the family of cytochrome P450 enzymes, including a ‘heme’ group speculated to act as the catalytic domain ^[25]^.

**Table 1:**
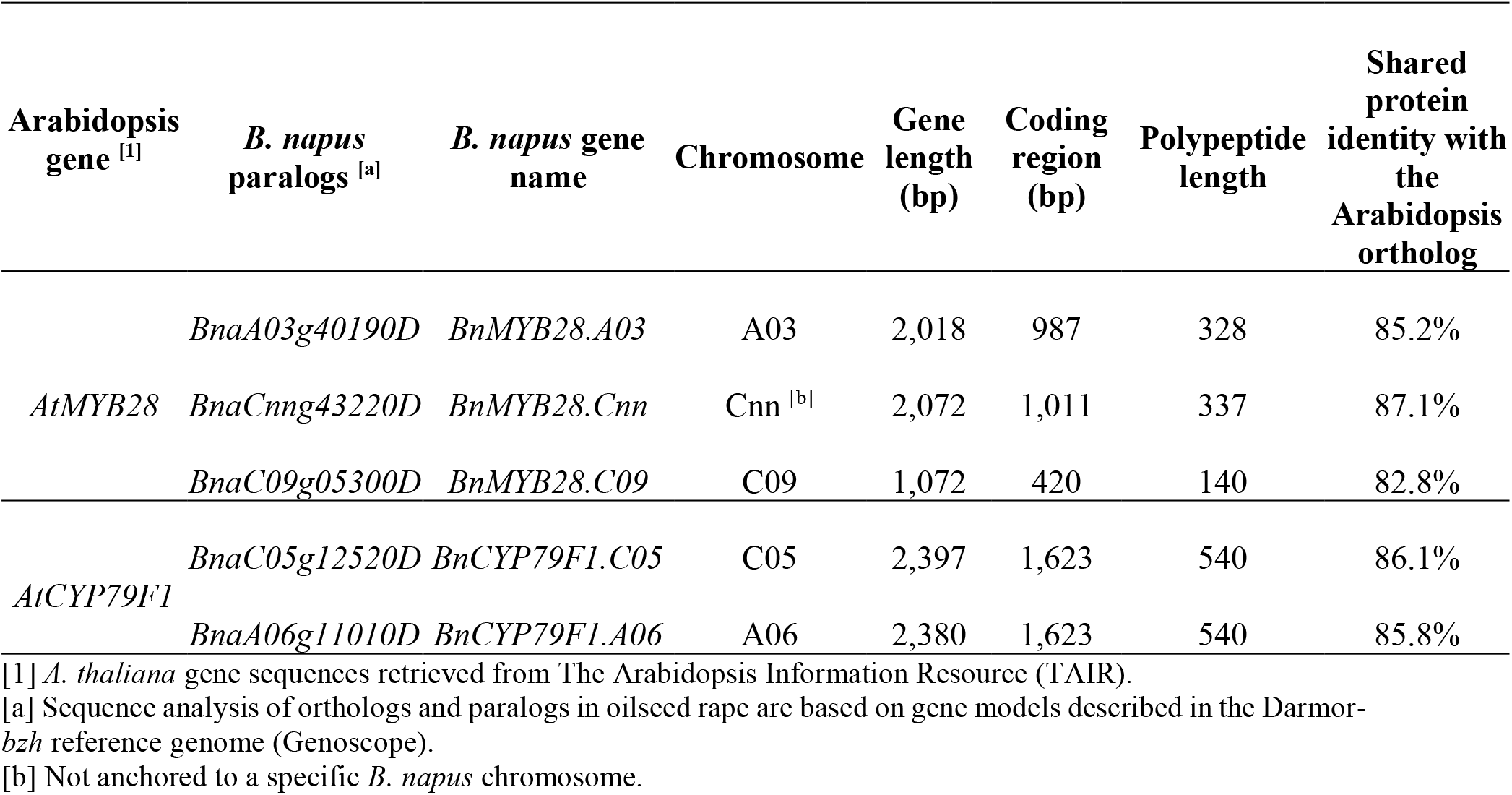
Features of *BnMYB28* and *BnCYP79F1* paralogs with homology to the Arabidopsis genes *AtMYB28* (AT5G61420) and *AtCYP79F1* (AT1G16410) in oilseed rape.

### Expression profiles of *BnMYB28* and *BnCYP79F1* genes reveal putative functional paralogs

For knock-out studies, we aimed to select highly expressed paralogs. Therefore, we investigated the expression profiles of *BnMYB28* and *BnCYP79F1* genes in the German winter-type inbred rapeseed Express617 by RT-qPCR. The relative expression of candidate genes was analyzed in leaves and seeds at growth stages 15, 25, 35, and 45 days after pollination (DAP).

The two *BnMYB28* paralogs *BnMYB28*.*C09* and *BnMYB28*.*A03* showed a thousand fold higher relative expression in leaves than seeds across all growth stages (Figure 1). *BnMYB28*.*C09* was the most highly expressed paralog in leaves. However, expression sharply declined 15 DAP and remained low during later stages of seed development (25-45 DAP).

**Figure 1:**
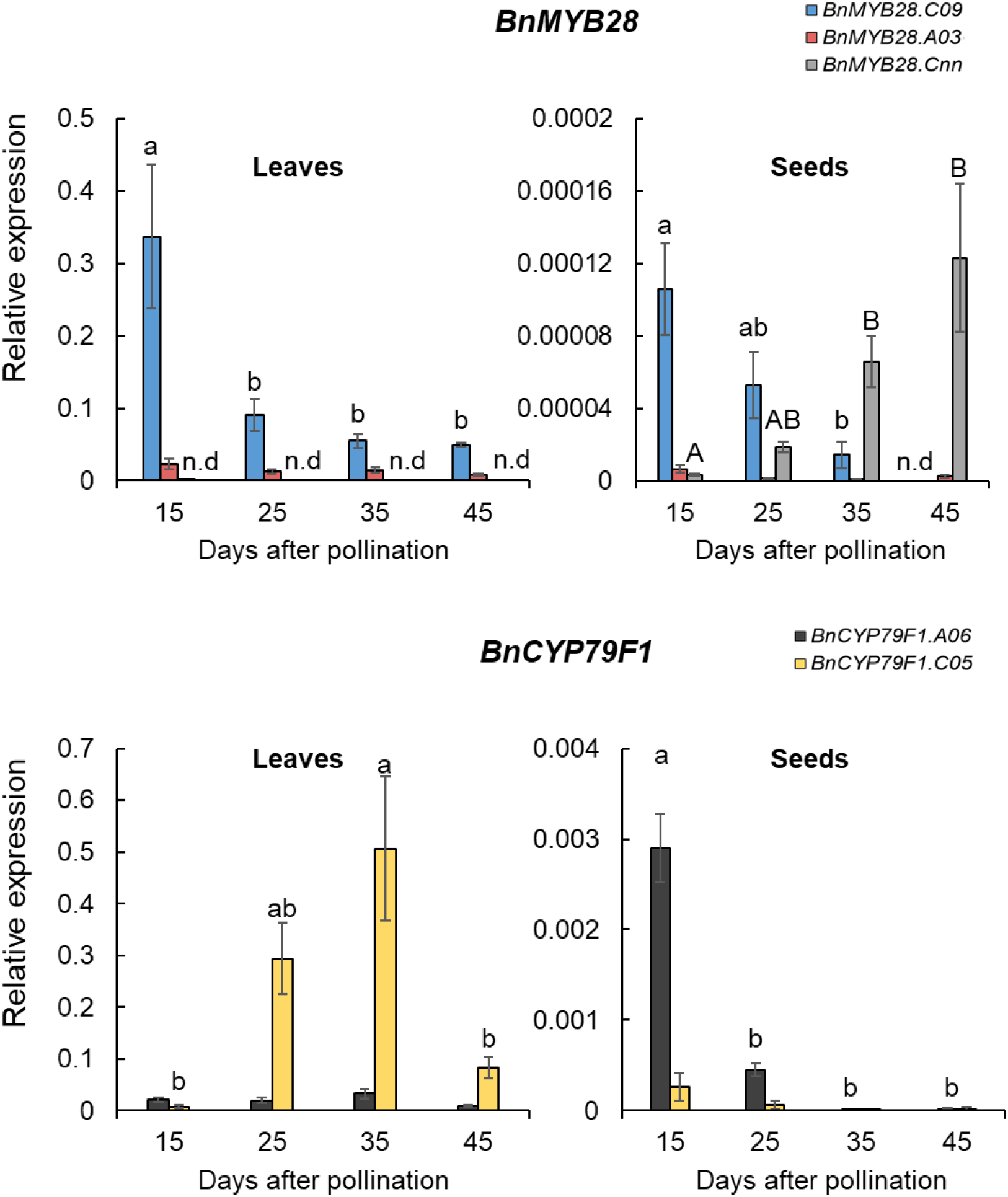
Relative expression of three *BnMYB28* and two *BnCYP79F1* paralogs in the winter-type oilseed rape Express617. Plants were grown in the greenhouse (20-25°C and 16 h light) after vernalization (4°C, 16 h light for 8 weeks). For each gene, leaves and seeds were sampled from five plants 15, 25, 35, and 45 days after pollination (five biological replicates). RT-qPCR was performed with three independent samples of each plant (three technical replicates) and the relative expression was calculated after normalization with two reference genes, *BnACTIN* and *BnGAPDH*. Error bars represent the standard error of the mean of five biological replicates, with three technical replicates each. Statistical significance was calculated with an ANOVA (*p*<0.05, linear model, grouping: Tukey test) using the R package “Agricolae”. Alphabets over bars represent statistical groups. n.d: not detectable.

The expression of *BnMYB28*.*A03* was consistently lower across all growth stages in the leaves, accounting for ∼17% of the expression levels of *BnMYB28*.*C09*. In leaves *BnMYB28*.*Cnn* expression was not detectable, whereas measurable expression was detected at later stages towards seed maturity. Conclusively, *BnMYB28*.*C09* and *BnMYB28*.*A03* were selected for further studies as putative functional paralogs because they were significantly expressed in the leaves.

Similar to *BnMYB28, BnCYP79F1* paralogs were significantly more expressed in the leaves than seeds, attaining a hundredfold higher expression level. There was also a significant difference between the two *BnCYP79F1* paralogs in leaves as the expression of *BnCYP79F1*.*C05* was more than tenfold higher than *BnCYP79F1*.*A06* (Figure 1). In contrast, the expression levels of *BnCYP79F1*.*A06* was higher than *BnCYP79F1*.*C05* in seeds. In the early stages of seed development (15 DAP), *BnCYP79F1*.*A06* was tenfold higher expressed than *BnCYP79F1*.*C05*, followed by a drastic decrease as the plants matured. In leaves, the expression of *BnCYP79F1*.*C05* sharply increased between 25 and 35 DAP and then decreased as the plants approached maturity. Interestingly, *BnMYB28* and *BnCYP79F1* genes displayed opposite expression patterns in the leaves. While *BnMYB28* was highly expressed at early stages (15 DAP) followed by a sharp decline, *BnCYP79F1* expression increased during later stages (25-35 DAP)

### EMS-induced mutants for selected *BnMYB28* and *BnCYP79F1* paralogs

We screened the M_2_ population for EMS-induced mutations in four genes (*BnMYB28*.*C09, BnMYB28*.*A03, BnCYP79F1*.*C05*, and *BnCYP79F1*.*A06*) using a conventional polyacrylamide gel-based assay. We detected 35 and 43 EMS-induced mutations in *BnMYB28* and *BnCYP79F1* paralogs, respectively (Supplementary Table 4), which could be classified into 6 nonsense, 50 missense, and 22 silent mutations (Figure 2). No splice site mutations were detected. Mutation frequencies ranged between 1/31.5-1/67.0 kb across the two gene families. On average, a frequency of one EMS-induced mutation per 43.6 kb was detected (Supplementary Table 5), which is in the range of estimations made by former studies on this EMS population ^[41; 42; 43; 44; 45; 46]^. Both *BnMYB28* nonsense mutations were located within the conserved DNA binding R2 domain. Additionally, seven missense mutations were also detected within the R2 domain. We found one nonsense mutation in the *BnCYP79F1*.*C05* paralog, while none could be detected for *BnCYP79F1*.*A06*. A missense mutation conferring a minor change to the protein folding due to the exchange of lysine with glutamic acid was selected. For further studies, we chose all M_3_ plants with nonsense mutations plus the *BnCYP79F1*.*A06* missense mutant. For ease of understanding, we assigned unique one-letter codes to wildtype and mutant alleles (Table 2).

**Figure 2:**
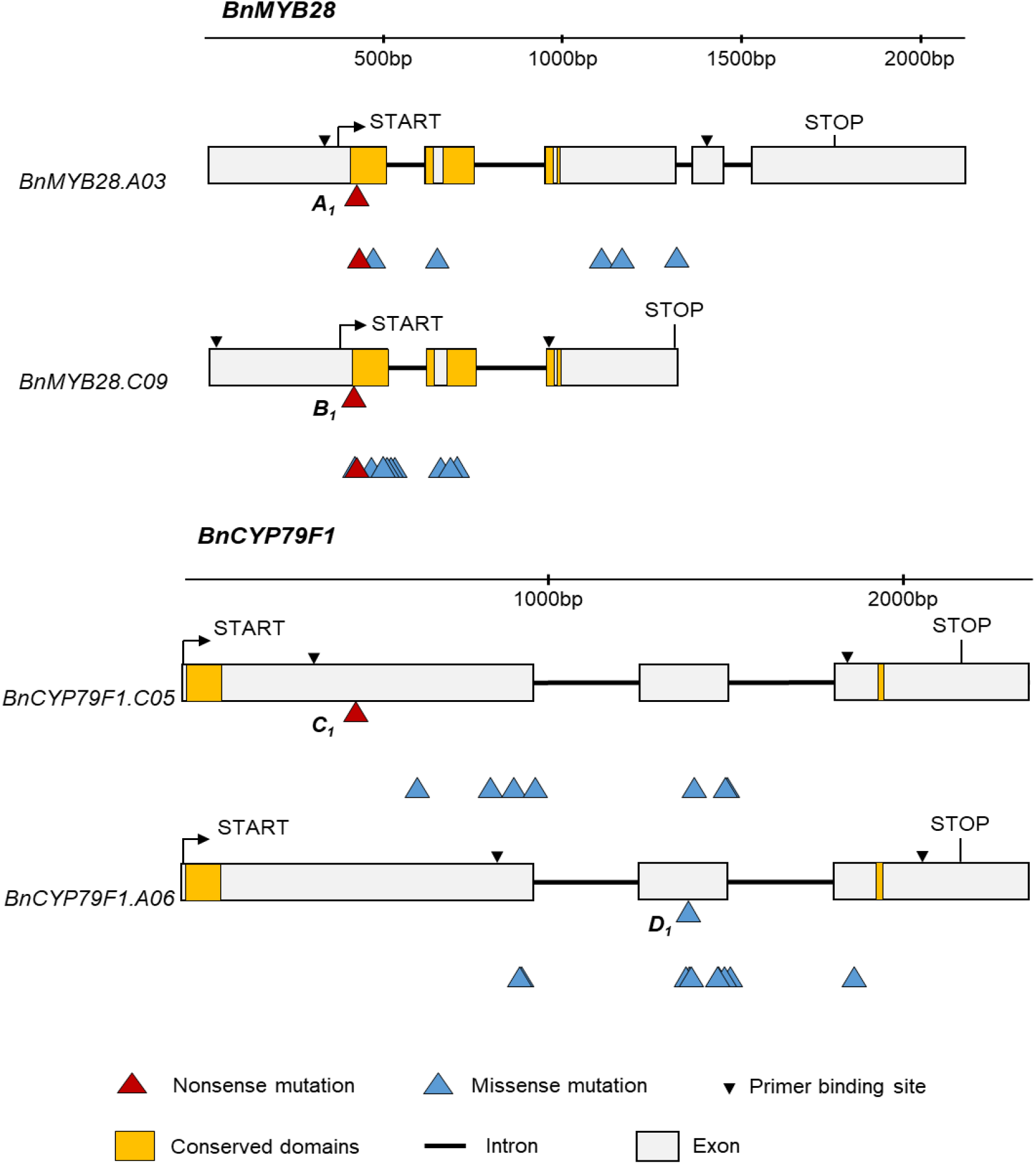
Structure of four *BnMYB28* and *BnCYP79F1* genes and EMS-induced nonsense and missense mutations. Allele identities are given next to the mutation site for those mutations used for further studies (refer to Table 2 for all allele codes). Regions coding for functional and conserved domains characteristic to the gene families are marked in orange boxes. START and STOP represent the translation start and stop sites, respectively. For *BnCYP79F1*, the 5’ untranslated regions are not defined on the Darmor-*bzh* reference genome.

**Table 2:**
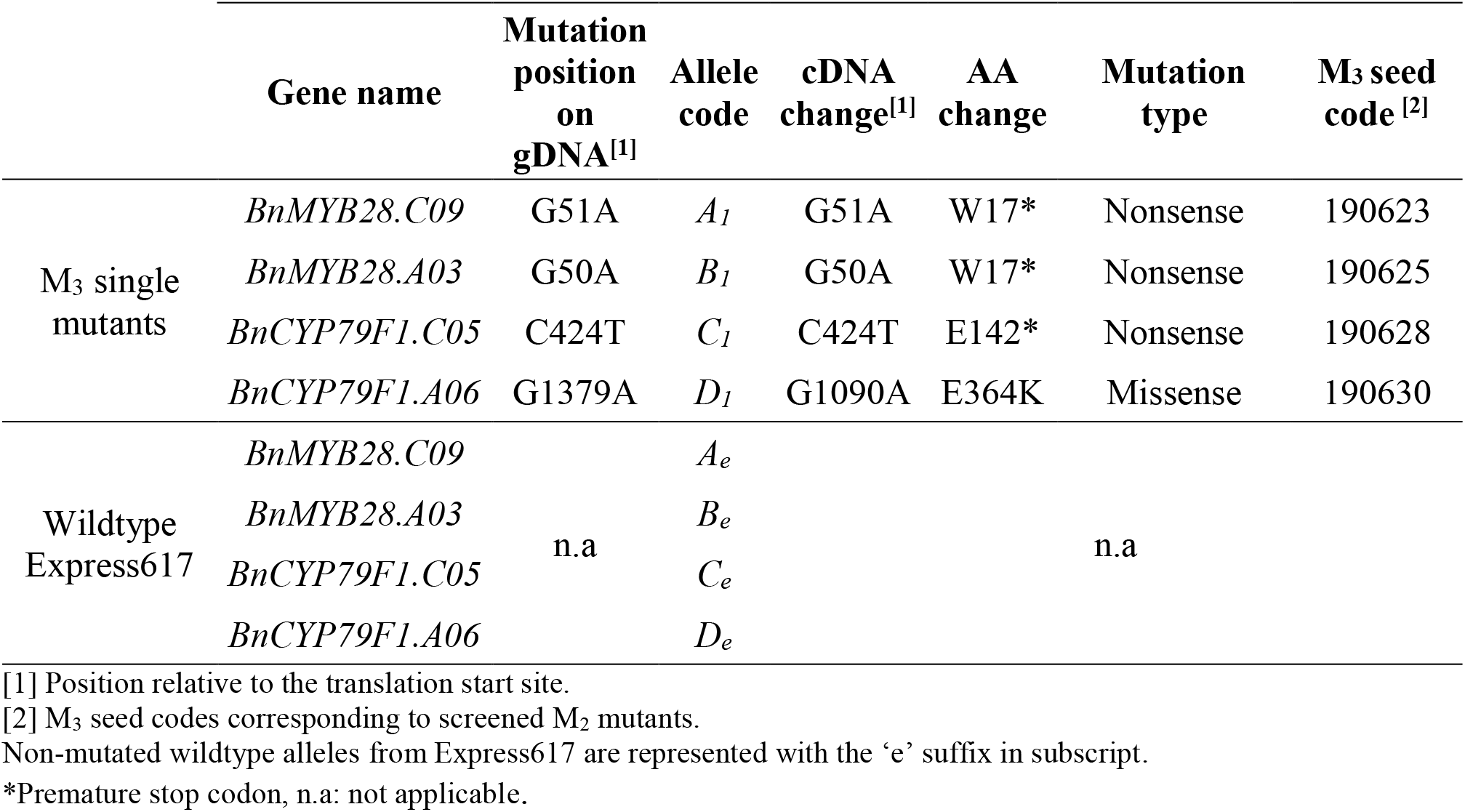
Allele codes assigned to EMS mutants and wildtype plants selected as crossing parents in this study. Single mutants were selected as crossing parents to combine single mutations for enhanced phenotypic effects. For each of the analyzed paralogs, mutants and wildtype parents were assigned unique allele codes.

Due to high functional redundancies of genes in the polyploid oilseed rape genome, single mutations rarely have a phenotypic effect. Therefore, we crossed single mutants of *BnMYB28*.*C09* and *BnMYB28*.*A03* and then separately for *BnCYP79F1*.*C05* and *BnCYP79F1*.*A06* to produce distinct double mutants of *BnMYB28* and *BnCYP79F1*, respectively. We genotyped M_3_ plants by generating PCR fragments encompassing the expected mutations (Supplementary Table 2) and confirmed single M_3_ mutants by Sanger sequencing of the PCR fragments (Supplementary Table 6). Mutant plants were selected for crossing experiments (Supplementary Table 7). *BnMYB28* and *BnCYP79F1* M_3_ single mutants were crossed with each other (referred to as ‘M_3_xM_3_’). F_1_ offspring were selfed to generate the F_2_ populations 200527 (Supplementary Figure 3A) and 200529 (Supplementary Figure 3B), respectively.

### Mutations in *BnMYB28* and *BnCYP79F1* confer a significant reduction in the aliphatic GSL content in seeds

We investigated the effect of EMS-induced mutations in *BnMYB28* and *BnCYP79F1* genes by quantitative and qualitative analysis of GSLs. Segregating individuals from two F_2_ populations (200527 and 200529) were analyzed (Supplementary Table 7) along with non-mutagenized Express617 plants as controls. F_2_ populations 200527 and 200529 originated from M_3_xM_3_ crosses of *BnMYB28* and *BnCYP79F1* single mutants, respectively.

First, F_2_ plants from the two populations were genotyped by Sanger sequencing. Four genotypes per population were selected for phenotypic studies. In F_2_ populations 200527 and 200529, plants homozygous for the wildtype alleles (*A*_*e*_*A*_*e*_*B*_*e*_*B*_*e*_ and *C*_*e*_*C*_*e*_*D*_*e*_*D*_*e*_), homozygous for one mutant allele (*A*_*1*_*A*_*1*_*B*_*e*_*B*_*e*_ or *A*_*e*_*A*_*e*_*B*_*1*_*B*_*1*_ and *C*_*1*_*C*_*1*_*D*_*e*_*D*_*e*_ or *C*_*e*_*C*_*e*_*D*_*1*_*D*_*1*_), and homozygous for two mutant alleles (*A*_*1*_*A*_*1*_*B*_*1*_*B*_*1*_ and *C*_*1*_*C*_*1*_*D*_*1*_*D*_*1*_) were selected for phenotyping (Supplementary Figure 3A and 3B).

In general, a significantly higher GSL content was observed in seeds than in leaves (Figure 3). In seeds of *BnMYB28* double mutants (F_2_ population 200527), the GSL content was significantly (*p* <0.05) reduced to 23.41 µmol/g DW compared to F_2_ plants homozygous for the wildtype alleles (47.28 µmol/g DW) and the non-mutagenized Express617 control plants (49.73 µmol/g DW), which corresponds to a significant GSL reduction by 50.5% and 52.9%, respectively (Figure 3). No significant differences were observed between the leaves of *BnMYB28* double mutants (0.96 µmol/g DW) and the Express617 control (0.86 µmol/g DW). Also, the *BnCYP79F1* double mutants (F_2_ population 200529) showed reduced seed GSL contents by 27.9% and 26.9% compared to the Express617 controls and the F_2_ plants homozygous for the wildtype alleles, respectively. However, the difference was not statistically significant (*p* <0.05). GSL contents in leaves varied between 0.9-1.5 µmol/g DW in the *BnCYP79F1* double mutants and 0.7-1.5 µmol/g DW in the wildtype F_2_ plants without statistically significant differences between the genotypes.

**Figure 3:**
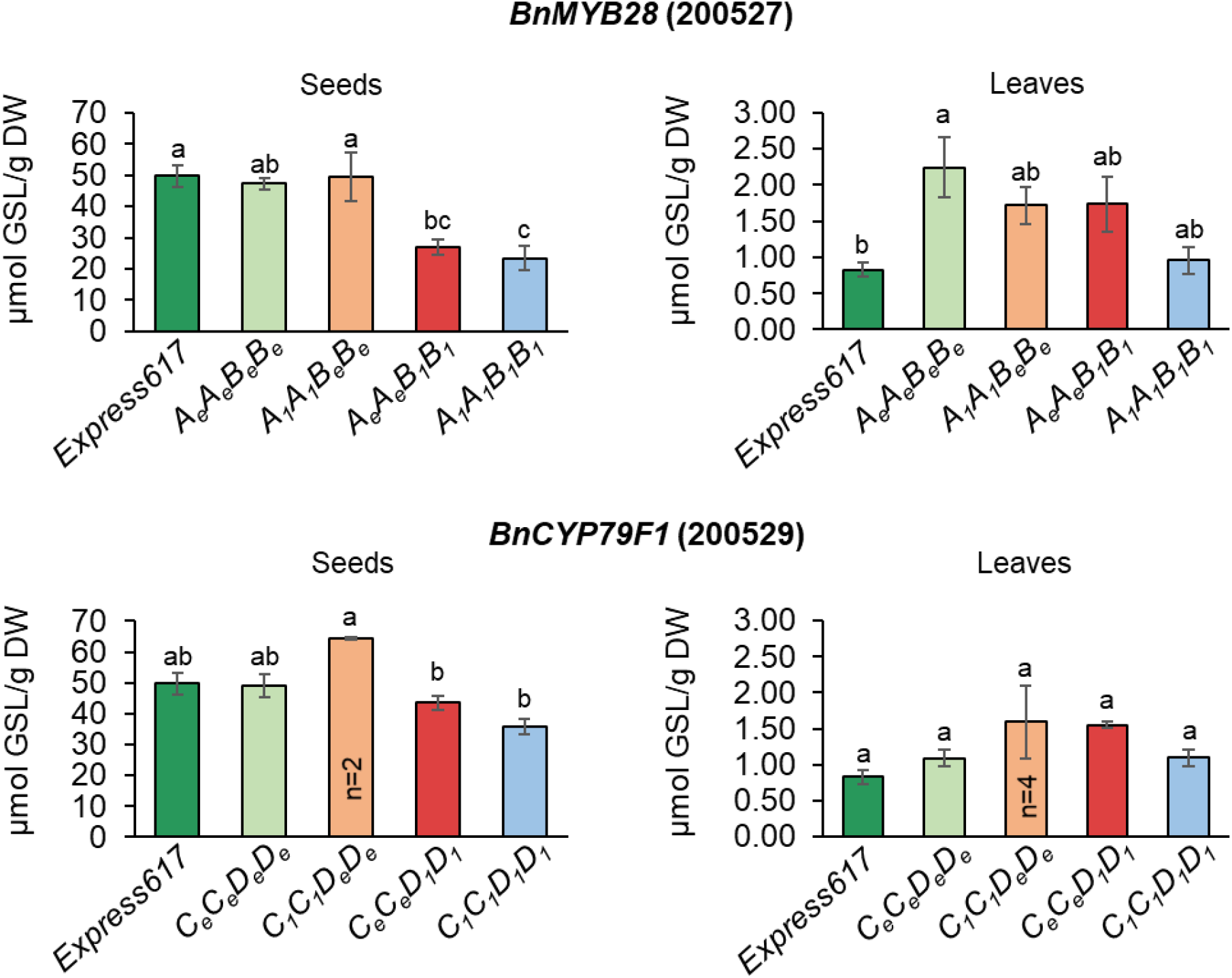
Seed and leaf glucosinolate contents in populations segregating for *BnMYB28* and *BnCYP79F1* mutations. F_2_ populations 200527 and 200529 segregating for *BnMYB28* and *BnCYP79F1* mutations, respectively, originated from M_3_xM_3_ crosses. Homozygous F_2_ double mutants (*A*_*1*_*A*_*1*_*B*_*1*_*B*_*1*_ and *C*_*1*_*C*_*1*_*D*_*1*_*D*_*1*_) were analyzed together with homozygous single mutants (*A*_*1*_*A*_*1*_*B*_*e*_*B*_*e*_, *A*_*e*_*A*_*e*_*B*_*1*_*B*_*1*,_ *C*_*1*_*C*_*1*_*D*_*e*_*D*_*e*,_ and *C*_*e*_*C*_*e*_*D*_*1*_*D*_*1*_), non-mutagenized Express617 and F_2_ plants homozygous for the wildtype alleles (*A*_*e*_*A*_*e*_*B*_*e*_*B*_*e*_ and *C*_*e*_*C*_*e*_*D*_*e*_*D*_*e*_). Leaf samples were taken 15 days after pollination and mature seeds (BBCH89) were used for glucosinolate determination. Error bars represent the standard error from five plants (n=5) per genotype with two exceptions mentioned in the figure. An ANOVA (*p <*0.05) was performed and the Tukey test (*p <* 0.05) was done for grouping. Different alphabets above error bars represent groups based on significance. All genotypes are as per designated allele codes given in Table 2.

Then, individual seed GSL profiles were analyzed in the same plants as performed for total GSL determinations. Single GSLs were identified by retention time and co-chromatography with commercial standards and quantified using individual calibration curves (Supplementary Table 3). Although the estimation of total GSL by summing up the major compounds identified by HPLC yielded generally higher values than the enzymatic method, the results did not show significant differences between mutants (Supplementary Figure 4). This suggested that our data evaluation for single GSL identification via HPLC and the sum of their concentrations were in line with the total GSL content estimated with the enzymatic test. Eight seed sample extracts were analyzed by LC-MS (Iontrap) to confirm the peak identity.

All previously calibrated glucosinolates were confirmed. Moreover, two additional aliphatic glucosinolates (gluconapoleiferin and glucoalyssin) and the major indolic glucosinolate 4-hydroxyglucobrassicin, for which no calibration standards were available, could be identified and were quantified based on their UV absorbance at 229 nm using sinigrin (as internal standard) and calibration factors from the literature ^[38]^.

In the seeds, we identified nine aliphatic GSLs (glucoiberin, progoitrin, epiprogoitrin, sinigrin, glucoraphanin, gluconapoleiferin, glucoalyssin, gluconapin, and glucobrassicanapin) in varying quantities and four other GSLs (4-hydroxyglucobrassicin, glucotropaeolin, glucobrassicin, and gluconasturtiin) in smaller amounts. In line with previous reports ^[9]^, the aliphatic GSL comprised the major portion (93.9%) of the seed GSL content in all genotypes studied here (Figure 4).

**Figure 4:**
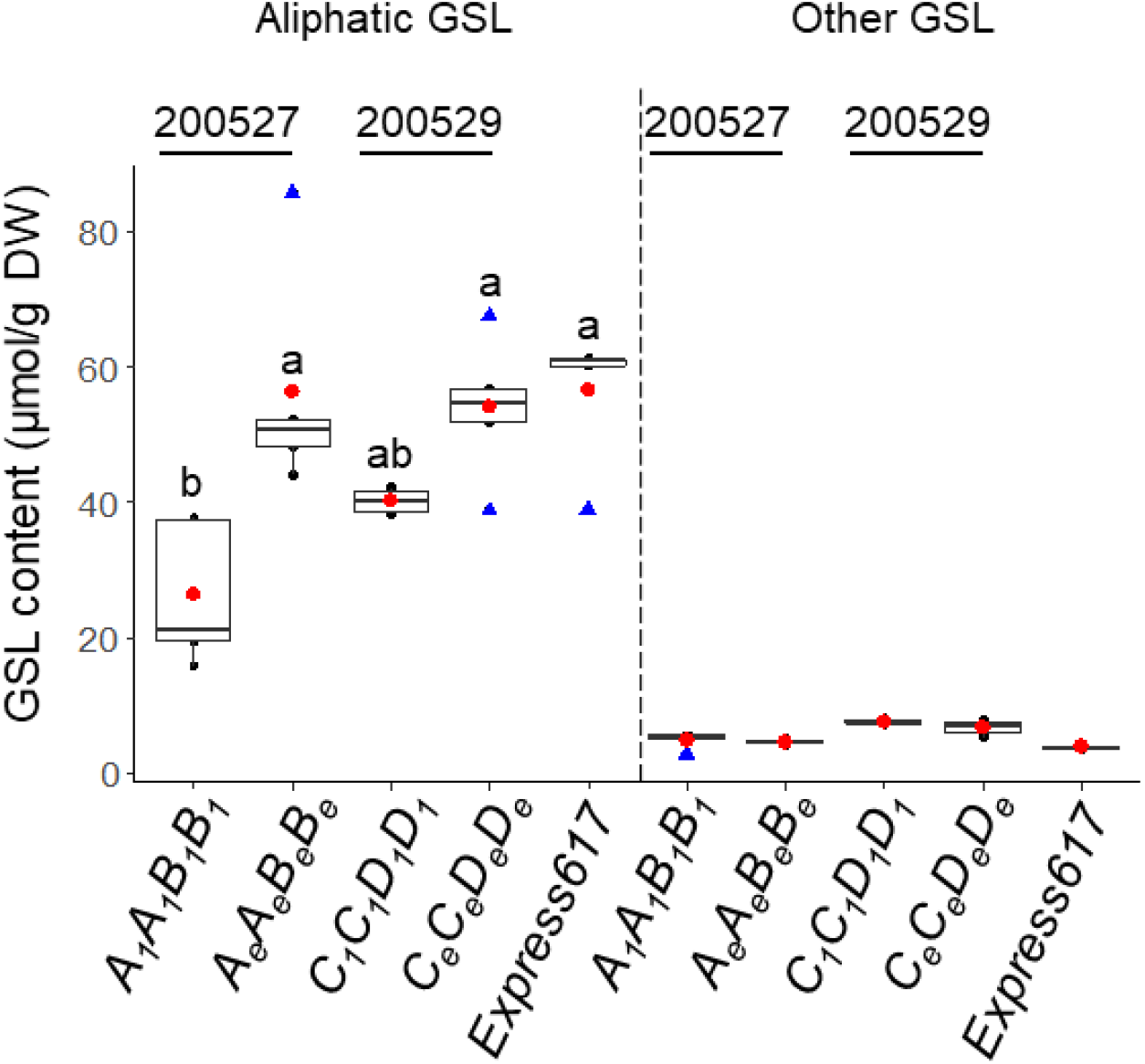
Analysis of major glucosinolate types in mature seeds of *BnMYB28* and *BnCYP79F1* mutants. Aliphatic and other GSL types measured from homozygous *BnMYB28* (genotype *A*_*1*_*A*_*1*_*B*_*1*_*B*_*1*_, seed code 200527) and *BnCYP79F1* (genotype *C*_*1*_*C*_*1*_*D*_*1*_*D*_*1*_, seed code 200529) originating from M_3_xM_3_ crosses. F_2_ plants homozygous for the wildtype alleles (*A*_*e*_*A*_*e*_*B*_*e*_*B*_*e*_ and *C*_*e*_*C*_*e*_*D*_*e*_*D*_*e*_) and non-mutated Express617 were used as controls. Plants were grown in the greenhouse. Mature seeds were harvested at BBCH89. GSL profiles for aliphatic and other GSL types (phenolic and indolic) were analyzed using HPLC. GSL content was calculated as µmol/g dry weight (DW). Individual and mean values are marked in black and red dots, respectively. Blue triangles represent outliers. Error bars represent the standard error of the mean from five biological replicates. An ANOVA test (*p <* 0.05) was performed and significant differences between groups were calculated by a Tukey test (*p <* 0.05). Different alphabets (a-d) above error bars represent groups based on significance. All genotypes are as per designated allele codes given in Table 2.

In *BnMYB28* F_2_ double mutants (population 200527), the progoitrin concentrations in seeds were 55.3% lower than in the Express617 controls (reduced from 36.32 µmol/g DW to 16.20 µmol/g DW) (Figure 5, Supplementary Table 8). The next abundant aliphatic compounds in the seeds were gluconapin and glucobrassicanapin. While glucobrassicanapin levels were drastically reduced by 87% from 5.26 µmol/g DW in the Express617 controls DW to 0.64 µmol/g DW in the double mutants, the gluconapin content was not significantly reduced. The minor aliphatic compound epiprogoitrin, whose synthesis starts from gluconapin, was reduced by 51% (0.57 µmol/g DW) in the double mutants compared to Express617 (1.17 µmol/g DW). The remaining seed GSLs analyzed did not exceed 3 µmol/g DW. Out of the 28 µmol/g DW reduction observed in the total seed GSL content of double mutants, the three major aliphatic GSLs accounted for 86% (24 µmol/g DW) of the total reduction. In the *BnCYP79F1* F_2_ double mutants, the progoitrin content was 32.4% lower than in Express617 (36.32 µmol/g DW compared to 24.5 µmol/g DW) (Figure 5, Supplementary Table 9). The glucobrassicanapin content was not altered in the *BnCYP79F1* double mutants. However, a 30.4% decrease in gluconapin content suggests that the *BnCYP79F1* mutations might have a bigger effect on the synthesis of short-chained 4C aliphatic than on the 5C aliphatic GSLs.

**Figure 5:**
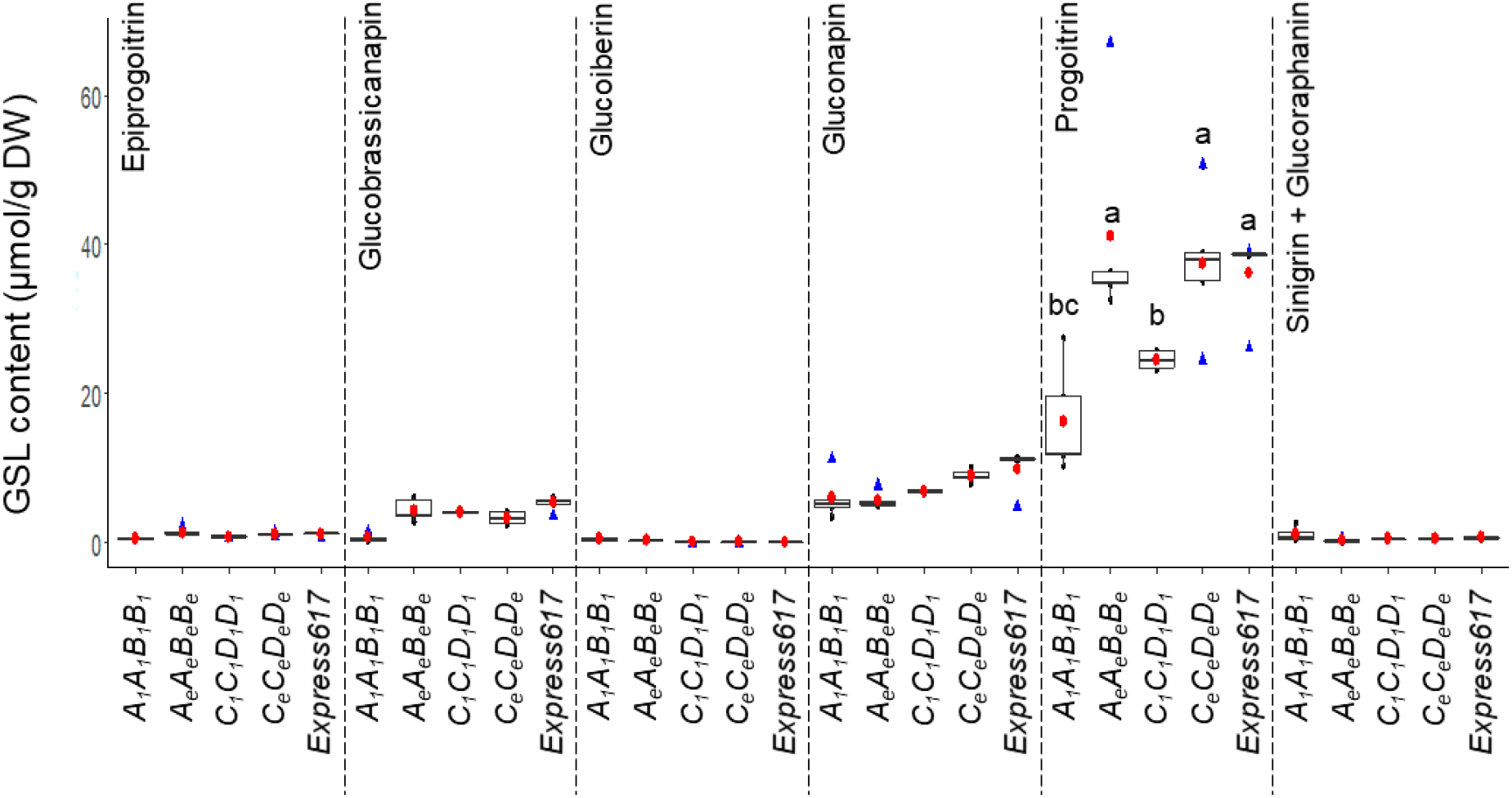
Analysis of aliphatic glucosinolates in mature seeds of *BnMYB28* and *BnCYP79F1* mutants. Individual aliphatic GSLs in homozygous F_2_ *BnMYB28* (genotype *A*_*1*_*A*_*1*_*B*_*1*_*B*_*1*_, seed code 200527) and *BnCYP79F1* (genotype *C*_*1*_*C*_*1*_*D*_*1*_*D*_*1*_, seed code 200529) double mutants originating from M_3_xM_3_ crosses. F_2_ plants homozygous for the wildtype alleles (*A*_*e*_*A*_*e*_*B*_*e*_*B*_*e*_ *and C*_*e*_*C*_*e*_*D*_*e*_*D*_*e*_) and non-mutated Express617 were used as controls. Aliphatic GSLs were identified and quantified using HPLC. The content was calculated as µmol/g dry weight (DW). Individual and mean values are marked in black and red dots, respectively. Blue triangles represent outliers. Error bars represent the standard error of the mean from biological replicates. An ANOVA test (*p <* 0.05) was performed and significant differences between groups were calculated by a Tukey test (*p <* 0.05). Different alphabets (a-e) above error bars represent groups based on significance. All genotypes are as per designated allele codes given in Table 2. n.d: not detectable.

## Discussion

Major anti-nutritive compounds like GSLs in the RSM pose a challenge for utilization as animal feed. Therefore, a major breeding goal is a reduction of the seed glucosinolate content (SGC) to an acceptable limit of <18 µmol/g dry weight. This study aimed to reduce the aliphatic GSL content in seeds by knocking out *BnMYB28* and *BnCYP79F1* genes involved in the biosynthesis of aliphatic GSLs in rapeseed. We demonstrate that independent knock-out mutants of the two genes possessed significantly reduced total and aliphatic GSLs, primarily progoitrin, in the seeds.

We targeted the aliphatic GSL biosynthesis pathway since the aliphatic profile comprises up to 92% of all GSLs reported from rapeseed ^[9]^. Moreover, major GSLs such as progoitrin that have adverse metabolic effects in animals belong to the aliphatic profile ^[47; 6; 16]^. We reasoned that functional mutations in genes involved in the secondary modification of GSLs might only confer an altered GSL profile and not a significant reduction in the overall content. Therefore, we selected *BnMYB28* and *BnCYP79F1* due to their prominent role in the core structure formation of aliphatic GSLs ^[25; 18]^. In *Arabidopsis*, a transcriptome study confirmed the role of sub-group 12 R2R3 MYB transcription factors in up-regulating almost all genes involved in the core structure formation of aliphatic GSLs ^[19]^. In associative transcriptomics and QTL mapping studies, former studies have found *MYB28* and *CYP79F1* to be strongly associated with a high aliphatic GSL content in rapeseed ^[48; 49]^. More recently, Kittipol et al. (2019) and Liu et al. (2020) have also identified *MYB28* as a significant gene controlling aliphatic GSL content in rapeseed using transcriptome and genome-wide association studies, respectively.

It has been demonstrated that the biosynthesis of GSLs occurs in vegetative parts, especially in rosette leaves and silique walls ^[50]^. Using histochemical analyses in *Arabidopsis*, Reintanz et al. (2001) have demonstrated that the activity of the biosynthesis gene *CYP79F1* is restricted to the silique walls, and almost untraceable expression levels were observed in the seeds. Moreover, *in silico* microarray analyses have shown that the expression of the *CYP79F1* and the *MYB28* transcription factors in *Arabidopsis* seeds was insignificant ^[51]^. The negligible expression levels of genes involved in the chain elongation and GSL core-structure formation steps in seeds strongly suggest their inability for the *de novo* synthesis of GSLs ^[51]^. Our expression analyses for *BnMYB28* and *BnCYP79F1* paralogs encompass seed setting and loading phases between 15-45 days after pollination (DAP). In our study, the expression profiles observed for the two biosynthesis genes complement previous studies since expression levels were significantly higher in leaves than seeds. *BnMYB28* paralogs were expressed thousand fold higher in the leaves. Relative expression levels increased during the early growth stages (15 DAP) with a gradual decrease as the plant approached maturity (45 DAP). This was expected since GSL biosynthesis increases as the plant transitions from the vegetative to the generative phase ^[52; 18]^. In *Arabidopsis*, Brown et al. (2003) have demonstrated that towards maturity, the reduction in leaf GSL content is concurrent with an increasing GSL content in the seeds. *BnMYB28*.*C09* showed the most significant expression levels over other gene copies in the leaves. In previous mapping studies, QTL significantly associated with high aliphatic GSL content in leaves and seeds of rapeseed were linked to *BnMYB28* on chromosome C09 ^[53; 29; 49; 54]^. Although *BnMYB28*.*Cnn* showed higher relative expression than other paralogs in seeds towards maturity, its levels were in trace amounts.

Moreover, its expression in leaves was undetectable. Since GSL biosynthesis is absent in seeds, we reason that the expression levels of *BnMYB28*.*Cnn* are too low to affect the seed GSL content. Most genes involved in aliphatic GSL biosynthesis, including *CYP79F1*, are under the transcriptional control of *MYB28* ^[19]^. This was evident since a significantly high relative expression of *BnMYB28*.*C09* at 15 DAP was followed by a significant increase in *BnCYP79F1*.*C05* expression levels in the leaves at 25 DAP. Based on these data, we reasoned paralogs *BnMYB28*.*C09* and *BnCYP79F1*.*C05* as our most promising candidates for functional analyses due to their significant expression levels in leaves, the primary site for GSL biosynthesis. However, due to the high functional redundancy in the polyploid rapeseed genome, we cannot wholly rely on single mutants of highly expressed paralogs for significant phenotypic effects. Therefore, we also considered *BnMYB28*.*A03* and *BnCYP79F1*.*A06* for pyramiding functional mutants for enhanced phenotypic effects.

Former studies demonstrating EMS-induced random mutagenesis in Brassicaceae crops have reported a wide range of mutation frequencies between 1/12 kb to 1/447 kb ^[55; 56; 57; 34; 58; 59]^. In this work, we screened the EMS-mutagenized population of the winter rapeseed ‘Express617’ developed by Harloff et al. (2012). Past studies on this resource have estimated varying mutation frequencies of 1/24 kb – 1/72 kb ^[41; 42; 43; 44; 45; 46]^. This variation is expected since frequency estimations depend on factors like the length of amplicons screened, the GC content within amplified fragments, and the number of pools screened for mutant detection. We estimated an average mutation frequency of 1/52.4 kb for *BnMYB28* and 1/34.7 kb for *BnCYP79F1*, which is well within the frequencies expected from this mutant population.

The functionality of *R2R3 MYB* transcription factors is determined by the DNA binding R2 and R3 domains and a nuclear localization signal ^[60; 18]^. We detected EMS-induced nonsense mutations within the conserved R2 domain in the first exons of *BnMYB28*.*C09* and *BnMYB28*.*A03*. Since both premature nonsense mutations were located in the R2 conserved domain, consequent transcripts are expected to lack the downstream R3 DNA binding and the vital NLS domains. For both single mutants, >95% of the resultant protein sequence is expected to be truncated. Therefore, we anticipate a complete loss-of-function of the *BnMYB28* transcription factor in selected mutants.

Within *BnCYP79F1*.*A06*, only missense mutations were found. A lack of nonsense mutations could be due to two reasons. First, the paralog-specific primers encompassed only 38% of the total cDNA sequence. Second, the amplicon possessed only five possible amino acid motifs with the possibility of converting to stop codons after the EMS-induced C→T or G→A transitions. Since we observed a nearly undetectable gene expression for *BnCYP79F1*.*A06* in the leaves compared to *BnCYP79F1*.*C05*, we speculate that its role in the biosynthesis process is less critical. Moreover, associative transcriptomics studies have reported *BnCYP79F1*.*C05* to be significantly correlated with the aliphatic GSL content in rapeseed ^[30]^.

Based on functional studies from *Arabidopsis* ^[25; 18]^ *and Brassica* crops ^[21; 22; 24; 27; 28]^, we expected that a knock-out of the two genes would severely influence the biosynthesis of short-chained aliphatic GSLs, especially progoitrin that accounts for ∼80% of all GSLs in the seeds ^[3]^. Since progoitrin is also the most abundant seed GSL type in rapeseed ^[9]^, we anticipated significant changes in the mutants. In this regard, a more significant effect from *BnMYB28* mutants was expected due to its central regulatory control over the aliphatic GSL biosynthesis ^[18; 19]^. Therefore, a knock-out of the *BnMYB28* is expected to downregulate several genes involved in the biosynthesis process. For validation, we suggest to analyze the expression of major downstream targets in *BnMYB28* double mutants, e.g., *MAM3* ^[61]^, which is involved in chain elongation and *CYP79F1, CYP79F2* ^[26]^, and *CYP83A1* ^[62]^ involved in the core structure formation. The mutants selected here are suitable for studying the role of *BnMYB28* as the ‘master regulator’ of the entire aliphatic GSL biosynthesis process in rapeseed.

Regarding *BnCYP79F1*.*A06*, only a missense mutation was available where a glutamic acid is replaced by a lysine. Therefore, a weaker phenotypic effect from the *BnCYP79F1* mutations can be explained by a putative compensation effect of *BnCYP79F1*.*A06*. Since *CYP79F1* can metabolize short and long-chained aliphatic GSLs in *Arabidopsis* ^[26]^, a possible sub-functionalization in the *B. napus* paralogs could explain the distinct functions of *BnCYP79F1*.*C05* and *BnCYP79F1*.*A06*. We speculate that *BnCYP79F1*.*A06* has a more significant impact on the C5 GSL metabolism than C4 aliphatic GSLs. This is because the sole knock-out of the *BnCYP79F1*.*C05* paralog in the *BnCYP79F1* double mutants resulted in significant reductions in the short-chained C4 aliphatic GSL progoitrin, whereas the C5 glucobrassicanapin content was not significantly reduced. For confirmation, functional analyses of *BnCYP79F1* knock-out mutants for both paralogs are needed.

It is known that the vegetative parts, especially the leaves are a major site for GSL biosynthesis. Although expression of biosynthesis genes is higher in vegetative parts, the seeds show a higher accumulation of GSLs in *Brassica* oilseeds since they are important sink tissues for GSLs in *Brassica* oil crops ^[63]^. Similarly, we observed a significantly higher GSL content in the seeds compared to the leaves. While this observation is in line with previous studies ^[52; 9; 51]^, it raises the question as to why GSL levels remain low and mostly unchanged in the leaves even in the mutants. Firstly, since the GSL content was already very low in the leaves, we did not observe any statistically significant reductions in the mutants. Secondly, we reason that the knock-out mutations in biosynthesis genes *BnMYB28* and *BnCYP79F1* described in this study confer restricted aliphatic GSL biosynthesis in the leaves. However, we speculate that a more significant phenotypic effect is realized in the seeds due to the subsequent activity of putative seed-specific GSL transporters ^[63; 64]^.

In conclusion, our study demonstrates the function of two major genes involved in the biosynthesis of aliphatic GSLs, the most abundant GSL class in rapeseed. Our results provide the first functional analysis by knock-out of *BnMYB28* and *BnCYP79F1* genes in rapeseed.

Mutants described in this study displayed significant reductions in the seed aliphatic GSL content that is well within commercial standards. In the future, investigating regulatory shifts in the complex GSL biosynthesis process and the seed-specific transport of GSLs could be crucial for achieving further GSL reduction. Our study provides a strong and promising basis for breeding rapeseed with improved meal quality in the future.

## Acknowledgments

We thank Monika Bruisch and Brigitte Neidhardt-Olf for their assistance in greenhouse experiments. We acknowledge the support provided by Jens Hermann and Prof. Dr. Wolfgang Bilger from the Department of Ecophysiology of Plants in Kiel for HPLC analytics. We also thank the Institute of Clinical Molecular Biology in Kiel for providing Sanger sequencing services. This work was financially supported by the Federal Ministry of Education and Research (BMBF) within the project IRFFA: Improved Rapeseed as Fish Feed in Aquaculture (grant number 031B0357B).

## Author Contributions

SJ, HJH and CJ designed the research. SJ conducted the experiments and analyzed the data. CW, MB and DT provided support for the qualitative assessment of glucosinolates. HJH and CJ supervised the research. SJ wrote the original draft. HJH, AA, MB, CW, DT and CJ reviewed and edited the manuscript. All authors participated in the discussion and revision of the manuscript. The authors read and approved the final manuscript.

## Data availability statement

The authors declare that data supporting the finding of this study are available from this manuscript and its supplementary information files. Extra data, information, and plant materials used/produced in this study are available from the corresponding author upon request.

