## Supplementary Figures for "Reduced glucosinolate content in oilseed rape (*Brassica napus* L.) by random mutagenesis of *BnMYB28* and *BnCYP79F1* genes"

Supplementary Figure 1

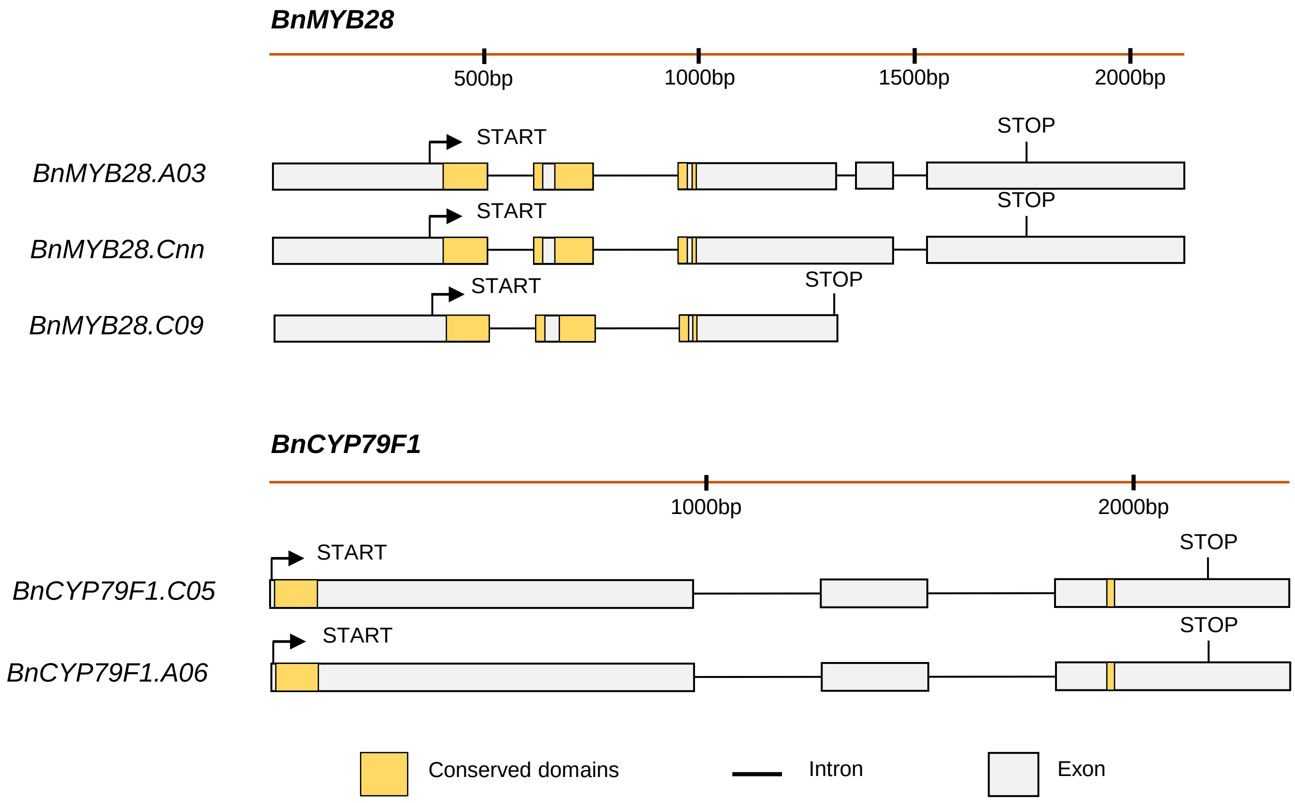


**Supplementary Figure 1:** Gene structures of three *BnMYB28* and two *BnCYP79F1* paralogs identified in rapeseed. START and STOP refer to the translation start and stop sites, respectively. Conserved domains characteristic of respective gene families are marked in yellow boxes. All gene models are based on the Darmor-*bzh* rapeseed reference genome. Paralog *BnMYB28.C09* was truncated but retained all conserved domains required for gene function. Information on the transcription start site for the *BnCYP79F1* paralogs is unavailable on the reference genome.

Supplementary Figure 2

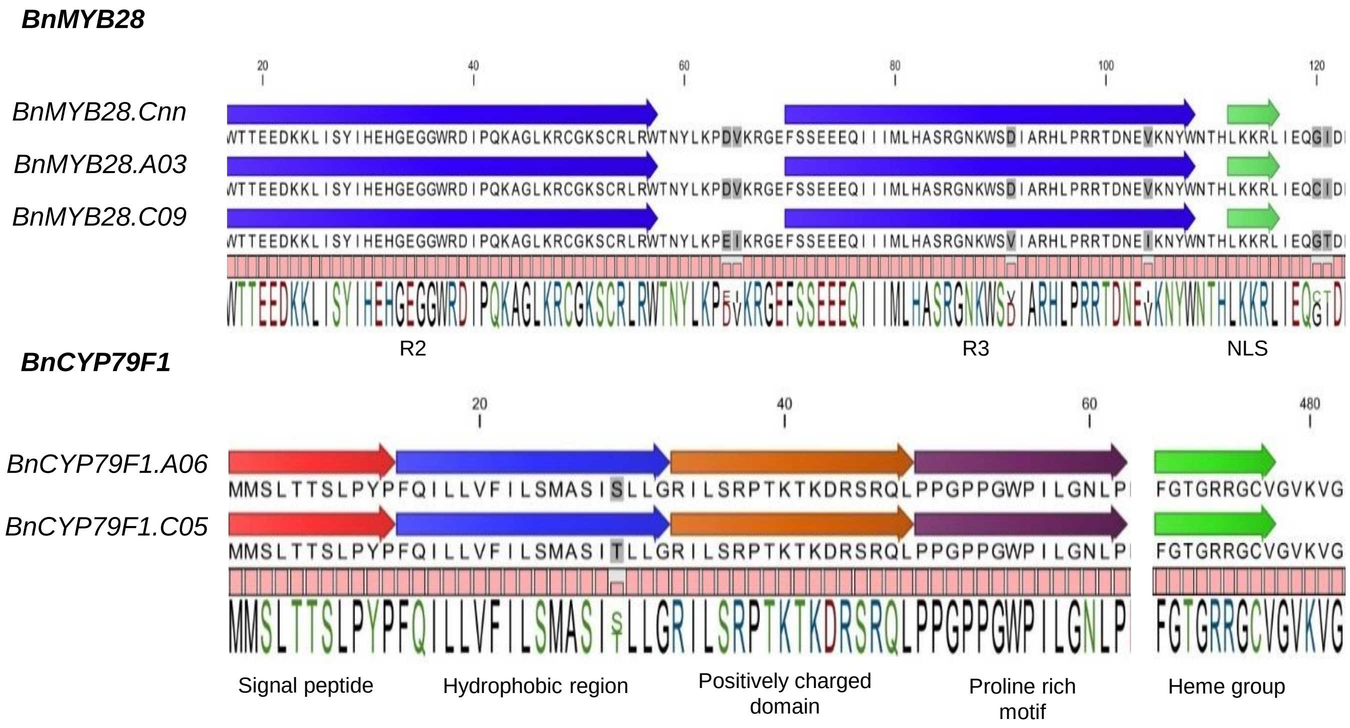


**Supplementary Figure 2:** Conserved domains observed in *MYB28* and *CYP79F1* paralogs of *B. napus.* The R2 and R3 DNA binding domains are highly conserved and characteristic of the MYB family transcription factors, followed by a nuclear localization signal (NLS) (Dubos et al., 2010). Five conserved domains have been reported from the family of cytochrome P450 enzymes. The heme group is speculated to act as a catalytic domain vital for enzyme function (Reintanz et al., 2001).

Supplementary Figure 3

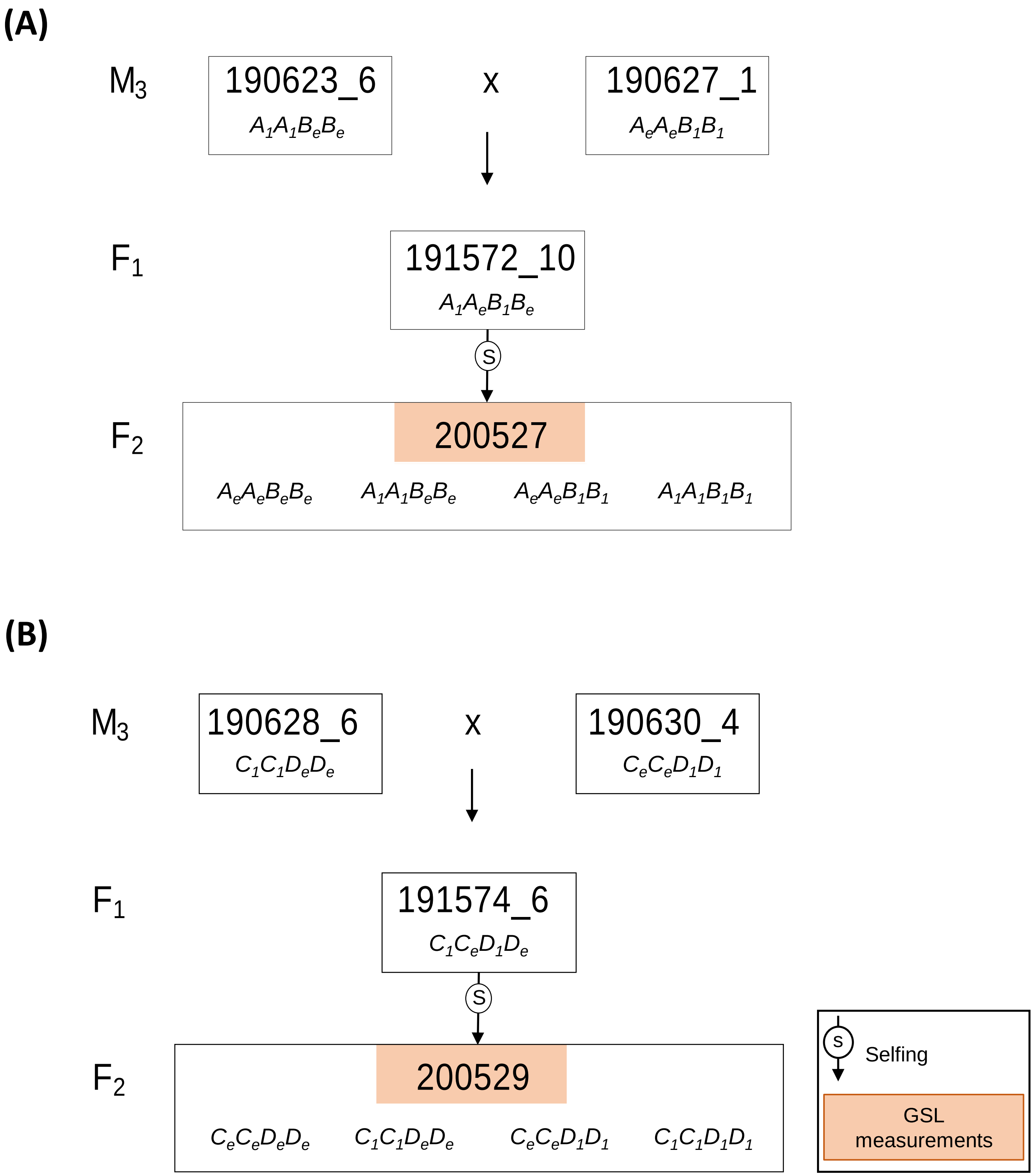


**Supplementary Figure 3:** Crossing scheme for producing double mutants of *BnMYB28* and *BnCYP79F1*. A) Crosses with *BnMYB28* $M_3$ single mutants to generate the $F_2$ population 200527. B) Crosses with *BnCYP79F1* $M_3$ single mutants to generate the $F_2$ population 200529. Plants with genotypes mentioned within boxes were selected in each generation for further experiments. The single-letter allele codes are shown in Table 2.

Supplementary Figure 4

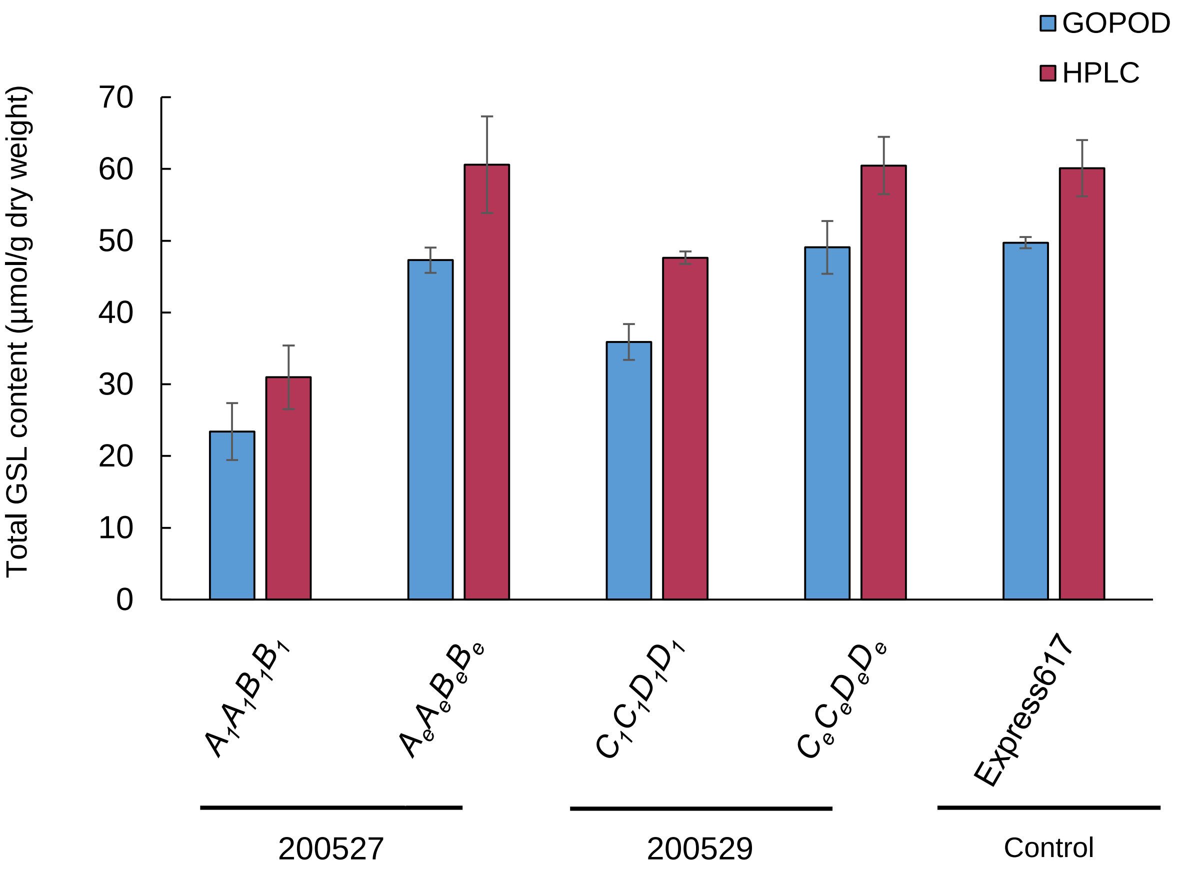


**Supplementary Figure 4:** Comparison of glucosinolate measurements by HPLC and the GOPOD enzymatic assay. The total GSL content determined by HPLC analyses is the sum of all individual peaks from chromatograms. Values were calculated after calibrating against GSL standards (Supplementary Table 3). The absolute GSL content was measured using the GOPOD enzymatic assay. *BnMYB28* and *BnCYP79F1* double mutants (genotypes $A_1A_1B_1B_1$ and $C_1C_1D_1D_1$) and wildtype plants ($A_eA_eB_eB_e$ and $C_eC_eD_eD_e$) segregating within the respective $F_2$ populations 200527 and 200529 originating from $M_3$x$M_3$ crosses were compared with the non-mutagenized Express617 control. For each of the analyzed genotypes, five biological replicates were analyzed. Error bars represent the standard error of the mean.
