## Supplementary Tables for "Reduced glucosinolate content in oilseed rape (*Brassica napus* L.) by random mutagenesis of *BnMYB28* and *BnCYP79F1* genes"

**Supplementary Table 1:** List of primers developed for RT-qPCR analyses.

| Gene name | Primer name | Primer sequence (5‘ → 3‘) | Tm (°C) | Amplicon length (bp) |
|---|---|---|---|---|
| *BnMYB28.C09* | GSL_rt_2_6_F | acagttcacgcctcttactcc | 60 | 144 |
| | GSL_2_6_R | gcttcatgcttctcgtgga | | |
| *BnMYB28.A03* | GSL_2_5_F | gcttcatgcttctcgtggt | 64 | 194 |
| | GSL_2_2_R | gcaaattctctggaggcgtgttgaca | | |
| *BnMYB28.Cnn* | GSL_2_3_F | cgaacagggtattgatccca | 58 | 264 |
| | GSL_rt_2_3_R | gtgaccttagccgcaactttg | | |
| *BnCYP79F1.C05* | GSL_3_1_F | cgccggaacacacgccatca | 64 | 258 |
| | GSL_rt_3_1_R | gcaaggaggttgtccgcttcaat | | |
| *BnCYP79F1.A06* | GSL_rt_3_2_F | ggtatacaaaccagagcgtcacctc | 64 | 222 |
| | GSL_3_2_R | cctctagacttaacggtccg | | |
| *BnACTIN2* | ACT1_F | tctggtgatggtgtgtctca | 60 | 141 |
| | ACT2_R | ggtcaacatgtaccctctctcg | | |
| *BnGAPDH* | GC1_F | ccgcttgcttcaacatcatt | 60 | 160 |
| | GC2_R | tctgaggtctttcgacgctg | | |

Primers were designed based on the Darmor-*bzh* reference genome (Genoscope).
F= Forward primer, R= Reverse primer

**Supplementary Table 2:** List of primers used for generating amplicons for mutant screening. One amplicon per paralog was used for mutant detection.

| | | Standard PCR [1] | | TILLING PCR [2] | | | |
|---|---|---|---|---|---|---|---|
| ***B. napus* gene name** | **Darmor ID** | **Primer name** | **Primer sequence (5‘ → 3‘)** | **TILLING primer name** | **Primer sequence (5‘ → 3‘)** | **TILLING amplicon length (bp) [3]** | **cDNA coverage [4]** |
| *BnMYB28.C09* | *BnaC09g05300D* | SJ_0082 | acagttcacgcctcttactcc | SJ_0094 | gagcttctctattctcatcctag | 902 | 65% |
| | | SJ_0083 | ggcttgtgagtcacgggatcag | SJ_0095 | gaccgaccacctaaagaccag | | |
| *BnMYB28.A03* | *BnaA03g40190D* | SJ_0084 | catgaaaacaccttgcagcta | SJ_0096 | gcattcttgggtgtttttgaggg | 1,706 | 73% |
| | | SJ_0085 | ctcgggattactaaccctgaaggc | SJ_0097 | gcgttggaactatcctcttc | | |
| *BnCYP79F1.C05* | *BnaC05g12520D* | SJ_0086 | gacaatgatgatgagccttacc | SJ_0098 | ccgacttcgcagaccggcct | 1,516 | 59% |
| | | SJ_0087 | cctctagacttaacggtcca | SJ_0099 | cggcctattccaggacgg | | |
| *BnCYP79F1.A06* | *BnaA06g11010D* | SJ_0088 | gttgtttggaaggagacatatc | SJ_0100 | cattgacgagagggtggagc | 1,180 | 38% |
| | | SJ_0089 | cctctagacttaacggtccg | SJ_0101 | ccatcatgatcgtcccgactttg | | |

[1] Standard PCR was done using unlabeled primers.

[2] TILLING PCR was done using a combination of unlabeled and infrared (IR) labeled primers. Forward and reverse primers were labeled with DY681 and DY781 probes, respectively. The TILLING PCR was done as a nested PCR using amplicons from the standard PCR as the template.

[3] Length of the amplicon analyzed for detection of EMS-induced mutations.

[4] Share of the total cDNA encompassed in the analyzed amplicon for detection of EMS-induced mutations.

**Supplementary Table 3:** HPLC analysis using commercial glucosinolate standards.

| Compound | Side chain | Retention Time (min) | HPLC calibration (linear regression) [a] |
|---|---|---|---|
| Glucoiberin | 3-me-sulfinylpropyl | 5.79 | (area-3367)/162966 |
| Progoitrin | (*2R*)-2 hydroxy-3-butenyl | 6.43 | (area+1631)/164513 |
| Epiprogoitrin | (*2S*)-2 hydroxy-3-butenyl | 6.60 | (area+18347)/157248 |
| Sinigrin | 2-propenyl | 6.66 | (area-11372)/199282 |
| Glucoraphanin | 4-me-sulfinylbutyl | 6.65 | (area+19466)/174317 |
| Gluconapin | 3-butenyl | 7.69 | (area-5783)/175044 |
| Glucobrassicanapin | 4-pentenyl | 8.77 | (area-9690)/162867 |
| Glucotropaeolin | benzyl | 8.91 | (area-9122)/198955 |
| Glucobrassicin | 3-Indolylmethyl | 9.40 | (area-11784)/552644 |
| Gluconasturtiin | phenethyl | 10.00 | (area-5888)/164863 |

[a] conversion from (area under the peak) to (nmol GSL)

**Supplementary Table 4**: All EMS-induced mutations detected in two *BnMYB28* and two *BnCYP79F1* paralogs. Mutation positions are relative to the translation start site.

| ***B. napus*** **gene name** | $M_2$ **plant name** [1] | $M_2$ **mutant zygosity** [2] | **Mutation position on gDNA** | **cDNA change** | **Amino acid change** | **Mutation type** |
|---|---|---|---|---|---|---|
| *BnMYB28.C09* | 70_F4 | Hom | G 12 C | G 12 C | K 4 N | Missense |
| | 70_H4 | Het | G 12 C | G 12 C | K 4 N | Missense |
| | 49_F2 | Het | G 26 A | G 26 A | G 9 E | Missense |
| | 66_B11 | Het | G 28 A | G 28 A | E 10 K | Missense |
| | 71_B4 | Hom | G 28 A | G 28 A | E 10 K | Missense |
| | 72_E9 | Het | G 31 A | G 31 A | G 11 R | Missense |
| | 71_E9 | Het | G 43 A | G 43 A | G 15 R | Missense |
| | 71_H9 | Hom | G 43 A | G 43 A | G 15 R | Missense |
| | 8_E5 | Het | G 43 A | G 43 A | G 43 R | Missense |
| | 64_C10 | Het | G 45 A | G 45 A | A 15 A | Silent |
| | 3_H2 | Het | G 51 A | G 51 A | W 17 * | Nonsense |
| | 11_H7 | Het | G 51 A | G 51 A | W 17 * | Nonsense |
| | 54_F10 | Het | C 78 T | C 78 T | I 26 I | Silent |
| | 43_E3 | Hom | G 97 A | G 97 A | G 33 R | Missense |
| | 54_C6 | Het | G 98 A | G 98 A | G 33 E | Missense |
| | 12_A9 | Hom | C 122 T | C 122 T | P 41 L | Missense |
| | 54_B9 | Het | G 133 A | G 133 A | G 45 R | Missense |
| | 54_C6 | Het | G 250 A | G 149 A | G 50 E | Missense |
| | 7_C6 | Het | C 289 T | C 188 T | P 63 L | Missense |
| | 7_D6 | Hom | C 289 T | C 188 T | P 63 L | Missense |
| | 76_D11 | Hom | G 303 A | G 202 A | G 68 S | Missense |
| | 64_A8 | Het | G 303 A | G 202 A | G 68 S | Missense |
| | 68_H6 | Hom | G 303 A | G 202 A | G 68 S | Missense |
| | 68_A11 | Het | G 323 A | G 222 A | E 74 E | Silent |
| | 45_B8 | Het | C 335 T | C 234 T | I 78 I | Silent |
| | 11_A5 | Het | G 329 A | G 329 A | Q 228 Q | Silent |
| *BnMYB28.A03* | 54_F3 | Hom | G 50 A | G 50 A | W 17 * | Nonsense |
| | 48_C8 | Hom | G 51 A | G 51 A | W 17 * | Nonsense |
| | 48_D8 | Hom | G 51 A | G 51 A | W 17 * | Nonsense |
| | 54_E1 | Het | G 61 A | G 61 A | E 21 K | Missense |
| | 48_H3 | Het | G 223 A | G 134 A | G 45 E | Missense |
| | 48_D3 | Hom | G 317 A | G 228 A | Q 76 Q | Silent |

| | | | | | | |
|---|---|---|---|---|---|---|
| | 50_E3 | Het | C 706 T | C 421 T | P 141 S | Missense |
| | 56_E12 | Hom | G 737 A | G 452 A | S 151 N | Missense |
| | 50_E3 | Het | C 899 T | C 614 T | T 205 I | Missense |
| *BnCYP79F1.C05* | 52_C8 | Het | C 424 T | C 424 T | E 142 * | Nonsense |
| | 64_C7 | Hom | C 560 T | C 560 T | T 187 I | Missense |
| | 64_F6 | Het | C 606 T | C 606 T | T 202 T | Silent |
| | 54_A6 | Het | C 654 T | C 654 T | F 218 F | Silent |
| | 53_B5 | Hom | G 770 A | G 770 A | G 257 D | Missense |
| | 63_E7 | Hom | G 850 A | G 850 A | E 284 K | Missense |
| | 65_E3 | Hom | C 956 T | C 956 T | P 319 L | Missense |
| | 65_A8 | Het | C 1399 T | C 1107 T | D 369 D | Silent |
| | 66_F6 | Het | G 1401 A | G 1109 A | R 370 K | Missense |
| | 53_C5 | Het | C 1430 T | C 1138 T | L 380 L | Silent |
| | 53_D5 | Het | C 1430 T | C 1138 T | L 380 L | Silent |
| | 56_A4 | Het | C 1430 T | C 1138 T | L 380 L | Silent |
| | 61_A10 | Het | C 1521 T | C 1229 T | T 410 I | Missense |
| | 61_C10 | Het | C 1521 T | C 1229 T | T 410 I | Missense |
| *BnCYP79F1.A06* | 61_A2 | Het | G 938 A | G 938 A | G 313 E | Missense |
| | 45_D7 | Het | G 1379 A | G 1090 A | E 364 K | Missense |
| | 41_A5 | Het | G 1384 A | G 1095 A | V 365 V | Silent |
| | 46_B6 | Het | G 1384 A | G 1095 A | V 365 V | Silent |
| | 53_B9 | Het | G 1385 A | G 1096 A | V 366 M | Missense |
| | 53_C9 | Het | G 1385 A | G 1096 A | V 366 M | Missense |
| | 46_F5 | Het | G 1394 A | G 1105 A | D 368 N | Missense |
| | 44_F12 | Het | G 1399 A | G 1110 A | R 370 R | Silent |
| | 44_H12 | Hom | G 1399 A | G 1110 A | R 370 R | Silent |
| | 44_G12 | Het | G 1399 A | G 1110 A | R 370 R | Silent |
| | 63_G10 | Het | C 1413 T | C 1124 T | S 375 F | Missense |
| | 69_F7 | Hom | C 1472 T | C 1183 T | P 395 S | Missense |
| | 74_A7 | Het | C 1472 T | C 1183 T | P 395 S | Missense |
| | 69_H7 | Het | C 1472 T | C 1183 T | P 395 S | Missense |
| | 43_F5 | Het | C 1473 T | C 1184 T | P 395 L | Missense |
| | 43_G5 | Het | C 1473 T | C 1184 T | P 395 L | Missense |
| | 48_A12 | Het | C 1473 T | C 1184 T | P 395 L | Missense |
| | 48_D12 | Het | C 1473 T | C 1184 T | P 395 L | Missense |
| | 76_A4 | Het | C 1473 T | C 1184 T | P 395 L | Missense |
| | 75_B9 | Het | C 1489 T | C 1200 T | V 400 V | Silent |
| | 42_B3 | Het | G 1511 A | G 1222 A | D 408 N | Missense |

| | | | | | |
|---|---|---|---|---|---|
| 61_A2 | Het | G 1511 A | G 1222 A | D 408 N | Missense |
| 51_C7 | Hom | C 1519 T | C 1230 T | T 410 T | Silent |
| 55_B2 | Hom | C 1520 T | C 1231 T | L 411 F | Missense |
| 63_E8 | Hom | C 1538 T | C 1249 T | P 417 S | Missense |
| 44_G6 | Het | G 1852 A | G 1289 A | G 430 D | Missense |
| 44_H12 | Hom | G 1898 A | G 1335 A | E 445 E | Silent |
| 44_F12 | Het | G 1898 A | G 1335 A | E 445 E | Silent |
| 44_G12 | Het | G 1898 A | G 1335 A | E 445 E | Silent |

[1] Name of the single $M_2$ mutant as per the Express617 EMS mutant resource (Harloff et al., 2012).
[2] Zygosity of EMS-induced mutations observed in single $M_2$ individuals.
Silent mutations do not confer changes in polypeptide sequences.
* Premature stop codon mutation.

**Supplementary Table 5:** Summary statistics of functional effects conferred by the EMS-induced mutations in *BnMYB28* and *BnCYP79F1* paralogs. One amplicon per paralog was used for screening EMS-induced mutations.

| | | | Number of detected mutations | | | | |
|---|---|---|---|---|---|---|---|
| **Gene name** | **cDNA coverage** | **Number of $M_2$ pools screened [1]** | **Nonsense mutations** | **Missense mutations** | **Silent mutations** | **Splice site mutations** | **Mutation frequency (1/kb) [2]** |
| *BnMYB28.C09* | 65% | 13 | 2 | 19 | 5 | 0 | 1/37.8 |
| *BnMYB28.A03* | 73% | 4 | 3 | 5 | 1 | 0 | 1/67.0 |
| *BnCYP79F1.C05* | 59% | 4 | 1 | 7 | 6 | 0 | 1/38.0 |
| *BnCYP79F1.A06* | 38% | 9 | 0 | 19 | 10 | 0 | 1/31.5 |
| | | | Total = 78 | | | | Average = 1/43.5 kb |

[1] The number of eight-fold (8x) two-dimensional (2-D) $M_2$ pools used for screenings.
[2] Calculated as the number of mutations per $M_1$ plant based on the number of analyzed $M_2$ families (8x 2D pools) and amplicon lengths.

**Supplementary Table 6:** Mutant genotyping in the $M_3$ generation to select crossing parents for the combination of single mutations.

| | | | | | | | | No. $M_3$ genotypes observed | | |
|---|---|---|---|---|---|---|---|---|---|---|
| **$M_3$ seed code** | **Gene name** | **$M_2$ genotype** | **Germination rate (%)** | **Mutation position on gDNA [1]** | **cDNA change** | **AA change [2]** | **Number of $M_3$ plants genotyped** | **Homozygous WT** | **Heterozygous mutant** | **Homozygous mutants** |
| 190623 | *BnMYB28.C09* | $A_1A_eB_eB_e$ | 80 | G 51 A | G 17 A | W 17 * | 12 | 6 | 4 | 2 |
| 190624 | *BnMYB28.C09* | $A_1A_eB_eB_e$ | 67 | G 51 A | G 17 A | W 17 * | 10 | 1 | 8 | 1 |
| 190625 | *BnMYB28.A03* | $A_eA_eB_1B_1$ | 60 | G 50 A | G 50 A | W 17 * | 4 | 0 | 0 | 4 |
| 190628 | *BnCYP79F1.C05* | $C_1C_eD_eD_e$ | 60 | C 424 T | C 424 T | E 142 * | 9 | 3 | 2 | 4 |
| 190630 | *BnCYP79F1.A06* | $C_eC_eD_1D_1$ | 80 | G 1379 A | G 1090 A | E 364 K | 11 | 0 | 0 | 11 |

Plants were genotyped using Sanger sequencing of PCR fragments encompassing expected EMS-induced mutations.
All genotypes have been named as per designated allele codes (Table 2).
[1] Position of EMS induced mutation on the gDNA from the translational start site
[2] Position of amino acid change on the polypeptide chain
*Premature stop codon

**Supplementary Table 7:** Genotypes of parents used for hand crosses and the offspring derived thereof to combine single EMS mutants of *BnMYB28* and *BnCYP79F1* and genotypes of selected crossing parents and their progenies used for phenotyping experiments.

| Crossing type | Parental genotypes | Genotype of selected $F_1$ progeny | $F_2$ seed code [a] | Genotypes used for phenotyping experiments [1] |
|---|---|---|---|---|
| $M_3$ x $M_3$ | $A_eA_eB_1B_1$ x $A_1A_1B_eB_e$ | $A_1A_eB_1B_e$ | 200527 | $A_eA_eB_eB_e$, $A_1A_1B_eB_e$, $A_eA_eB_1B_1$, $A_1A_1B_1B_1$ |
| | $C_eC_eD_1D_1$ x $C_1C_1D_eD_e$ | $C_1C_eD_1D_e$ | 200529 | $C_eC_eD_eD_e$, $C_1C_1D_eD_e$, $C_eC_eD_1D_1$, $C_1C_1D_1D_1$ |

All genotypes have been named as per designated allele codes (refer to Table 2).
Non-mutated wildtype alleles from Express617 are represented by the 'e' suffix in subscript.
[1] For quantitative and qualitative glucosinolate determination.
[a] Segregating $F_2$ population originating from heterozygous *BnMYB28* and *BnCYP79F1* double mutants (originating from direct $M_3$x$M_3$ crosses).

**Supplementary Table 8:** Summary statistics of individual glucosinolates identified with corresponding concentrations estimated in seeds of *BnMYB28* EMS mutants and controls. Analyses were done using high-performance liquid chromatography.

| | **$F_2$ population [1]** | | | | | |
|---|---|---|---|---|---|---|
| | Population 200527 | | | | Control | |
| **Glucosinolate type [a]** | *BnMYB28* _DM | SD | *BnMYB28* _WT | SD | Express617 | SD |
| Glucoiberin | 0.53 | 0.25 | 0.30 | 0.08 | 0.10 | 0.01 |
| Progoitrin | 16.20 | 6.52 | 41.22 | 13.05 | 36.32 | 5.09 |
| Epiprogoitrin | 0.57 | 0.16 | 1.37 | 0.41 | 1.18 | 0.27 |
| Sinigrin + Glucoraphanin | 1.12 | 0.86 | 0.27 | 0.21 | 0.64 | 0.08 |
| Gluconapoleiferin | 0.04 | 0.02 | 0.11 | 0.07 | n.d | |
| Glucoallysin | 1.03 | 0.41 | 2.82 | 0.72 | 2.94 | 0.05 |
| Gluconapin | 6.05 | 2.70 | 5.66 | 1.06 | 9.93 | 2.52 |
| 4-Hydroxyglucobrassicin | 2.56 | 0.61 | 2.31 | 0.16 | 2.23 | 0.32 |
| Glucobrassicanapin | 0.64 | 0.48 | 4.33 | 1.30 | 5.27 | 0.88 |
| Glucotropaeolin | 0.75 | 0.14 | 0.76 | 0.06 | 0.50 | 0.08 |
| Glucobrassicin | 0.26 | 0.12 | 0.13 | 0.03 | 0.20 | 0.36 |
| Gluconasturtiin | 1.08 | 0.24 | 1.07 | 0.13 | 0.73 | 0.16 |
| **Sum (µmol/g DW)** | **30.83** | | **60.35** | | **60.04** | |

[1] Experiments with plants originating from direct $M_3xM_3$ crosses without backcrossing.

[a] Estimated based on comparison of retention time and co-chromatography with commercial standards. GSL concentrations were calculated as µmol/g dry weight of tissue analyzed (five biological replicates per genotype). *BnMYB28*_DM: *BnMYB28* double mutant (genotype $A_1A_1B_1B_1$), *BnMYB28*_WT: Plants with wildtype alleles for *BnMYB28* segregating within the same $F_2$ population, Express617: non-mutagenized parent

n.d: Not detectable SD: Standard deviation

**Supplementary Table 9:** Summary statistics of individual glucosinolates identified with corresponding concentrations estimated in seeds of *BnCYP79F1* EMS mutants and controls. Analyses were done using high-performance liquid chromatography.

| | **$F_2$ population [1]** | | | | | |
|---|---|---|---|---|---|---|
| | Population 200529 | | | | Controls | |
| **Glucosinolate type [a]** | *BnCYP79F1*_DM | SD | *BnCYP79F1*_WT | SD | Express617 | SD |
| Glucoiberin | 0.01 | 0.01 | 0.04 | 0.02 | 0.10 | 0.01 |
| Progoitrin | 24.55 | 1.09 | 37.53 | 8.41 | 36.32 | 5.09 |
| Epiprogoitrin | 0.78 | 0.07 | 1.12 | 0.17 | 1.18 | 0.27 |
| Sinigrin + Glucoraphanin | 0.45 | 0.07 | 0.42 | 0.03 | 0.64 | 0.08 |
| Gluconapoleiferin | 0.08 | 0.02 | 0.05 | 0.02 | n.d | |
| Glucoallysin | 3.28 | 0.15 | 2.47 | 0.55 | 2.94 | 0.05 |
| Gluconapin | 6.91 | 0.20 | 8.91 | 0.73 | 9.93 | 2.52 |
| 4-Hydroxyglucobrassicin | 4.40 | 0.19 | 3.81 | 0.68 | 2.23 | 0.32 |
| Glucobrassicanapin | 4.02 | 0.15 | 3.33 | 0.68 | 5.27 | 0.88 |
| Glucotropaeolin | 1.02 | 0.04 | 0.90 | 0.13 | 0.50 | 0.08 |
| Glucobrassicin | 0.31 | 0.05 | 0.19 | 0.06 | 0.20 | 0.36 |
| Gluconasturtiin | 1.62 | 0.07 | 1.56 | 0.13 | 0.73 | 0.16 |
| **Sum (µmol/g DW)** | **47.43** | | **60.33** | | **60.04** | |

[1] Experiments with plants originating from direct $M_3$x$M_3$ crosses without backcrossing.

[a] Estimated based on comparison of retention time and co-chromatography with commercial standards. GSL concentrations were calculated as µmol/g dry weight of tissue analyzed (five biological replicates per genotype).

*BnCYP79F1*_DM: *BnMYB28* double mutant (genotype $C_1C_1D_1D_1$), *BnCYP79F1*_WT: Plants with wildtype alleles for *BnCYP79F1* segregating within the same $F_2$ population, Express617: non-mutagenized parent

n.d: Not detectable SD: Standard deviation
